# The ELIXIR Core Data Resources: fundamental infrastructure for the life sciences

**DOI:** 10.1101/598318

**Authors:** Rachel Drysdale, Charles E. Cook, Robert Petryszak, Vivienne Baillie-Gerritsen, Mary Barlow, Elisabeth Gasteiger, Franziska Gruhl, Jürgen Haas, Jerry Lanfear, Rodrigo Lopez, Nicole Redaschi, Heinz Stockinger, Daniel Teixeira, Aravind Venkatesan, Niklas Blomberg, Christine Durinx, Johanna McEntyre, ELIXIR Core Data Resource Forum

## Abstract

**Motivation:** Life science research in academia, industry, agriculture, and the health sector depends critically on free and open data resources. ELIXIR (www.elixir-europe.org), the European Research Infrastructure for life sciences data, has identified a set of Core Data Resources within Europe that are of most fundamental importance for the long-term preservation of biological data. We explore characteristics of their usage, impact and assured funding horizon to assess their value and importance as an infrastructure, to understand sustainability of the infrastructure, and to demonstrate a model for assessing Core Data Resources worldwide.

**Results:** The nineteen resources currently designated ELIXIR Core Data Resources form a data infrastructure in Europe which is a subset of the worldwide open life science data infrastructure. We show that, from 2014 to 2018, data managed by the Core Data Resources more than tripled while staff numbers increased by less than a tenth. Additionally, support for the Core Data Resources is precarious: together they have assured funding for less than a third of current staff after four years.

Our findings demonstrate the importance of the ELIXIR Core Data Resources as repositories for research data and knowledge, while also demonstrating the uncertain nature of the funding environment for this infrastructure. ELIXIR is working towards longer-term support for the Core Data Resources and, through the Global Biodata Coalition, aims to ensure support for the worldwide life science data resource infrastructure of which the ELIXIR Core Data Resources are a subset.

**Supplementary information:** Supplementary data are available at *Bioinformatics* online.

## 1 Introduction

Life science data resources have been used extensively in academia and industry for well over two decades and are increasingly used in clinical settings. These resources are critical for ensuring the reproducibility and integrity of the entire life sciences research enterprise (Bourne *et al.*, 2015). Despite their importance, many are supported in whole or in part by short-term grants, and there is little coordination of funding across these resources (Berman, 2008; Gabella *et al.*, 2017; https://www.biorxiv.org/content/10.1101/110825v3).

ELIXIR (www.elixir-europe.org) brings together life sciences resources from across Europe. More than 20 countries contribute to ELIXIR’s infrastructure with scientific tools and databases, as well as compute infrastructure, standards for interoperability, and training. Here, we focus on existing, well-established data resources. One of ELIXIR’s goals is to support the most valuable, used and useful resources, i.e., those with a very high scientific impact. To fulfill this goal ELIXIR has created a formal process to identify the most critical life sciences data resources in Europe, designated ELIXIR Core Data Resources (https://www.elixir-europe.org/platforms/data/core-data-resources; Durinx *et al.*, 2016). There are currently 19 Core Data Resources (CDRs, Table 1), spanning a broad range of life sciences data types including genes and genomes, proteins, chemistry, molecular structures and interactions, and the research literature. The process to identify these resources (Durinx *et al.*, 2016) uses a set of qualitative and quantitative indicators of scientific and technical quality and impact. The indicators fall into five categories: Scientific focus and quality of science; Community served by the resource; Quality of service; Legal and funding infrastructure, and governance; Impact and translational stories. The resources identified in this way are of fundamental importance to the wider life sciences community and the longterm preservation of biological data: they are comprehensive, are considered an authority in their fields, are of high scientific quality and provide a high level of service delivery. It is of critical importance that these resources are sustained for the benefit of all researchers.

**Table 1:** List of ELIXIR’s Core Data Resources

| Resource Name | Overview | Reference |
| --- | --- | --- |
| ArrayExpress | Data from high-throughput functional genomics experiments | Athar *et al.*, 2019 |
| BRENDA | Database of enzyme and enzyme-ligand information | Jeske *et al.*, 2019 |
| CATH | Hierarchical domain classification of protein structures PDB | Sillitoe *et al.*, 2019 |
| ChEBI | Dictionary of molecular entities focused on ‘small’ chemical compounds | Hastings *et al.*, 2016 |
| ChEMBL | Database of bioactive drug-like small molecules | Mendez *et al.*, 2019 |
| EGA | Personally identifiable genetic and phenotypic data | Lappalainen *et al.*, 2015 |
| ENA | Nucleotide sequencing information | Harrison, 2019 |
| Ensembl | Genome browser for vertebrate genomes | Cunningham, *et al.*, 2019 |
| Ensembl Genomes | Genome browser for non-vertebrate genomes, with sites for bacteria, protists, fungi, plants, and invertebrate Metazoa | Kersey *et al.* 2018 |
| Europe PMC | Repository to life sciences articles, books, patents and clinical guidelines | Levchenko *et al.*, 2018 |
| Human Protein Atlas | Information on human protein-coding genes | Uhlén *et al.*, 2015 |
| IMEx Consortium<br>(IntAct and MINT) | IntAct: experimentally-verified molecular interactions<br>MINT: experimentally-verified protein-protein interactions | Orchard *et al.*, 2012 |
| InterPro | Functional analysis of protein sequences | Mitchel *et al.*, 2019 |
| Orphadata | Comprehensive, high-quality datasets related to rare diseases | Rath *et al.*, 2012 |
| PDBe | Biological macromolecular structures | Mir *et al.*, 2018 |
| PRIDE | Mass spectrometry-based proteomics data | Perez-Riverol *et al.*, 2019 |
| SILVA | Resource for quality checked and aligned ribosomal RNA sequence data | Glöckner *et al.*, 2017 |
| STRING | Known and predicted protein-protein interactions. | Szklarczyk *et al.*, 2019 |
| UniProt | Comprehensive resource for protein sequence and annotation data | UniProt Consortium 2019 |

Here we characterise the Core Data Resources using a subset of the indicators helpful for portraying aspects of the utility and value of the resources to the research community over time. Rather than considering data resources individually, we discuss the Core Data Resources as a collective entity, that together form an integrated life sciences data infrastructure. As previously described (Durinx *et al.*, 2016), managers of the Core Data Resources supply indicator data as part of the selection process, with updates provided on an annual basis. We have for the first time used data collected from the Core Data Resources covering the years 2013-2018, to characterise this emerging infrastructure as a whole. We present summaries of their use and impact, their interconnectedness as an ecosystem, and describe their context within the Open Data and FAIR landscape. In this way we demonstrate their foundational role within the life sciences, which contrasts with the short-term nature of the assured funding horizon under which they operate.

## 2 Methods

Qualitative and quantitative information to support the life cycle management of the Core Data Resources is gathered by a defined and iterative process that has been described elsewhere (Durinx *et al.*, 2016, and https://zenodo.org/record/1194123#.XG_anC10eL5). This work depends on close collaboration between the managers of the ELIXIR Core Data Resources, ELIXIR, and tools and infrastructure providers who facilitate access to the necessary information.

Data were collected in two phases. For the first round of Core Data Resource selection (https://f1000research.com/documents/7-1711) a Case Document was prepared by the applicant resource managers, providing information about 23 indicators (Durinx *et al.*, 2016) for the calendar years 2013-2015. Annual updates were subsequently requested for 2016-2018 from the selected Core Data Resources. For the second round of selection (https://f1000research.com/documents/7-1712) the applicants provided indicator data for the calendar years 2014-2016, later updated with 2017 and 2018 data.

In the following section, the methods used to generate each Figure are described in turn. The data from which the Figures were generated, and additional specific descriptions of methodology and techniques, can be found in the accompanying Supplementary Data.

Figure 1: *Data entries:* This indicator corresponds to Indicator 3b “Data entries - Total, cumulative” from Durinx *et al.*, 2016. Each CDR decides which data entity is its primary entry type and provides counts on an annual basis. Data types include nucleic acid and protein sequences, genomes and metagenomes, macromolecular structures, molecular complexes, publications, complex assemblies, and articles from the scientific literature. The items that constitute “Data Entries” therefore vary between the resources, but the counts down the years are of the same entity for each CDR.

**Figure 1.**
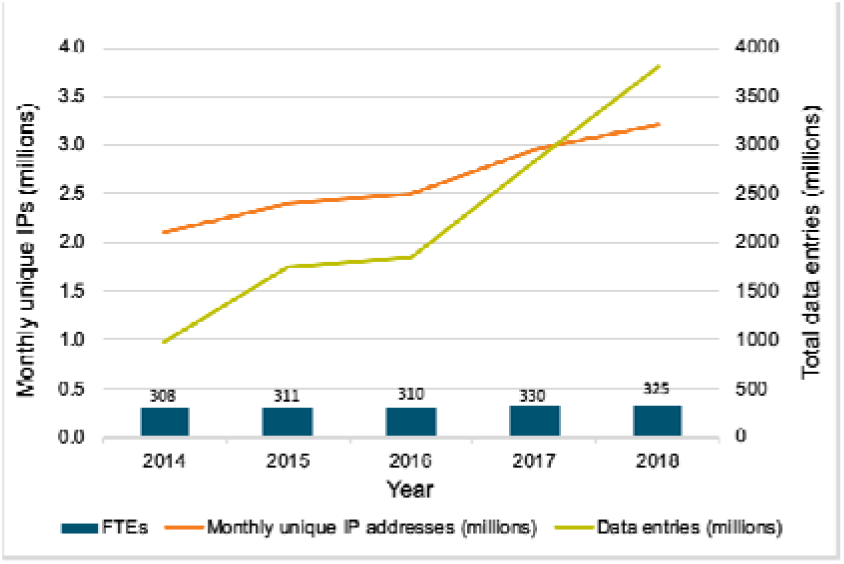
Scale of the Core Data Resources.

### Users

This indicator corresponds to Indicator 2a “Overall usage: visitors” from Durinx *et al.*, 2016. The CDRs are, by virtue of the selection criteria, open to all users with no requirement to register for an account. Because usage is unrestricted, determining the number of users poses a challenge. One way to measure the user community is to count the average monthly web access for each year in terms of unique IP addresses (Metwally and Paduano, 2011; Metwally *et al.*, 2014). This is necessarily a proxy for user numbers and both under- and over-reporting is possible, e.g., users may access resources from multiple devices and thus have multiple IP addresses, and users may also be connected using systems with dynamic IP address assignment: both situations generate more IP addresses than individuals. Conversely, some institutions representing hundreds or thousands of users may appear as a single IP address, leading to underreporting. Additionally, a single IP address that accesses different Core Data Resources will be counted separately for each resource. On balance, the number of unique IP addresses is almost certainly an overestimate of the number of users.

Web access can be measured with web analytics or log analytics. Web analytics (“web page tagging”) is based on tags that are embedded in web pages and cookies stored on a user’s device, and are typically collected through services such as Google Analytics. Log analytics are based on the analysis of IP address data collected on the server hosting the resource. Although web analytics are generally easier to set up, they do not track 100 percent of requests because JavaScript may not be executed on the client side, for example when cookies or image downloading are blocked, as is typical on mobile devices. Log analytics, on the other hand, are more complicated to set up, requiring dedicated hardware and infrastructure. The system used depends on the technology that is preferred by the hosting institution of the respective CDRs. For 14 CDRs, the estimation of the usage was based on log analytics, and for five resources on Google Analytics. When both measures were reported, log analytics figures were chosen for this analysis.

### Staff effort in Full Time Equivalents (FTEs)

This corresponds to Indicator 1d “Staff effort: number of FTEs per year for the past 2–3 years” from Durinx *et al.*, 2016 and includes curators, bioinformaticians and technical staff representative of each calendar year as reported by each resource manager. This reflects the staff required to develop and maintain a data resource. The distribution of types of staff varies between the CDRs. In Deposition Databases, such as ArrayExpress or ENA, the focus is on technical staff and bioinformaticians. By contrast, knowledgebases, for example the Human Protein Atlas or UniProt, add layers of value through teams of highly qualified curators who manually analyse and standardise research data. Each resource uses its own method to settle on an FTE count to provide in its annual update, then uses that same method for each year. This consolidates part-time and full-time contributors to the equivalent number of full-time positions, so it does not necessarily reflect the actual number of people involved in the resource. It is likely that the FTE count recorded for CDRs housed within large bioinformatics institutes underestimates the actual staff effort required to support such resources, due to economies of scale and institutional support provided within those large institutes. By contrast, a resource operating in a smaller institute may be the only hosted service, and must depend on local staff for all computational infrastructure management.

Figure 2: *Literature mentions and citations:* This corresponds to Indicator 2c “Usage in research as measured through citation in the literature” from Durinx *et al.*, 2016. This indicator aims to evaluate how the CDRs contribute to specific research projects. For each CDR three different types of citation indicators in Europe PMC have been counted on a yearly basis: a) mentions of the name of the CDR, through mining of the patterns of the resource name, b) citation of individual accessions from each CDR, identified through mining of the patterns of their unique identifiers, and c) citations of selected Key Articles describing the individual resources in other publications (see Supplementary Data for further details). In Figure 2, mentions of the CDR by name and citations of individual accessions are combined, while citations to key articles are shown as a separate column. We note that citations of key articles were counted individually for each CDR. These citations were not checked for duplications so the cumulative citation count will over-count publications that cite key articles from two or more CDRs.

**Figure 2.**
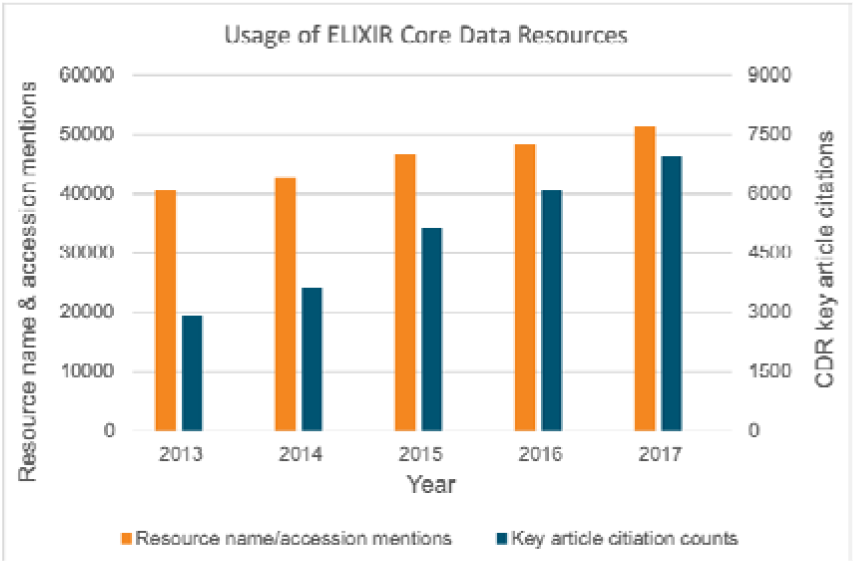
Usage of ELIXIR Core Data Resources in research.

These citation indicators conservatively estimate usage of CDRs in research projects as the estimates are constrained by the number of full text papers available in Europe PMC, *de facto* excluding the non-open access literature. Mining resource-name mentions was carried out for 16 of the 19 CDRs: BRENDA, SILVA and Orphadata were not included in the initial list of CDRs, and have not yet been folded into the “Resource Name Mentions” text mining pipeline. Mining of entry identifiers was carried out for 13 of those 16 resources: three resources do not assign their own unique identifiers to individual data sets (see Supplementary Data for further details). A caveat to this methodology is that use of certain resources has become such a routine element of everyday research practice that they are rarely cited. This is the case for literature repositories such as Europe PMC, which are heavily used but rarely explicitly acknowledged. Additionally, while initiatives to encourage data citation are gaining traction (https://doi.org/10.25490/a97f-egyk), these are relatively recent and not yet comprehensively adopted. These factors contribute to significant, but difficult-to-quantify, undercounting of literature citations to the CDRs.

Figure 3: *Categories of the top 20 CDR-citing journals*: This is related to Indicator 2c “Usage in research as measured through citation in the literature” from Durinx *et al.*, 2016. Three citation indicators of CDRs were collected: a) mentions of the name of the CDR, through mining of the patterns of the resource name, b) citation of individual records within the CDR, through mining of the patterns of their unique identifiers, and c) citations of selected Key Articles describing the individual resources in other publications (see Supplementary Data for details - this data was collected on 15th August 2018). For each unique PMID across the three citation indicators, the journal title and citation count were retrieved from Europe PMC. The top 20 CDR-citing journals were identified and mapped to a set of categories, based on the category model used in the <u>Scimago Journal & Country Rank</u> (https://www.scimagojr.com/journalrank.php). Finally, the number of citations to CDRs in all three indicators in journals within each category were tallied and plotted against the categories. Citation counts are cumulative from the publication date for each article to 15th August 2018, when the data were collected.

**Figure 3.**
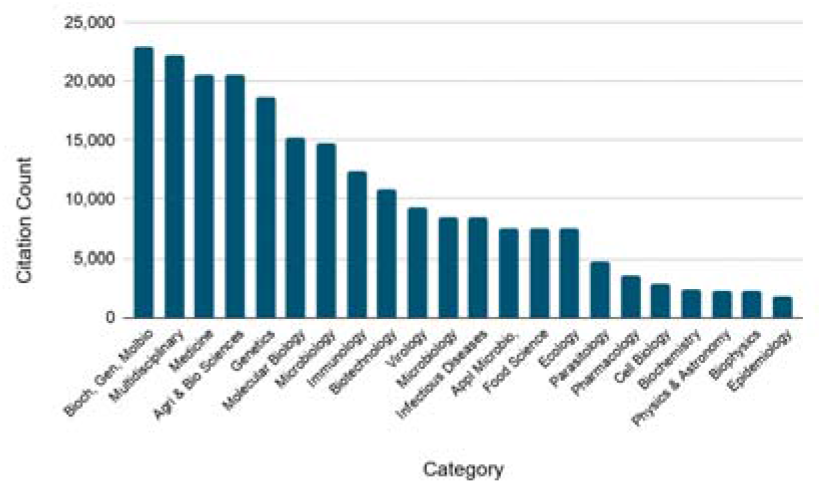
Cumulative citation counts, to 15th August 2018, for the categories of scientific fields in which the 20 journals that most frequently cite the Core Data Resources are active.

Figure 4: *Core Data Resource interconnectivity:* This is related to Indicator 2d “Dependency of other resources” from Durinx *et al.*, 2016. Lists of the data resources with which each CDR directly exchanges data, generally using application programming interface calls, were requested from the CDR managers. In Figure 4 the number of individual types of data exchanged by each pair of CDRs is plotted. The relationships are expressed in a chord diagram, with the arc width weighted according to the number of links from each CDR to the other CDRs.

**Figure 4.**
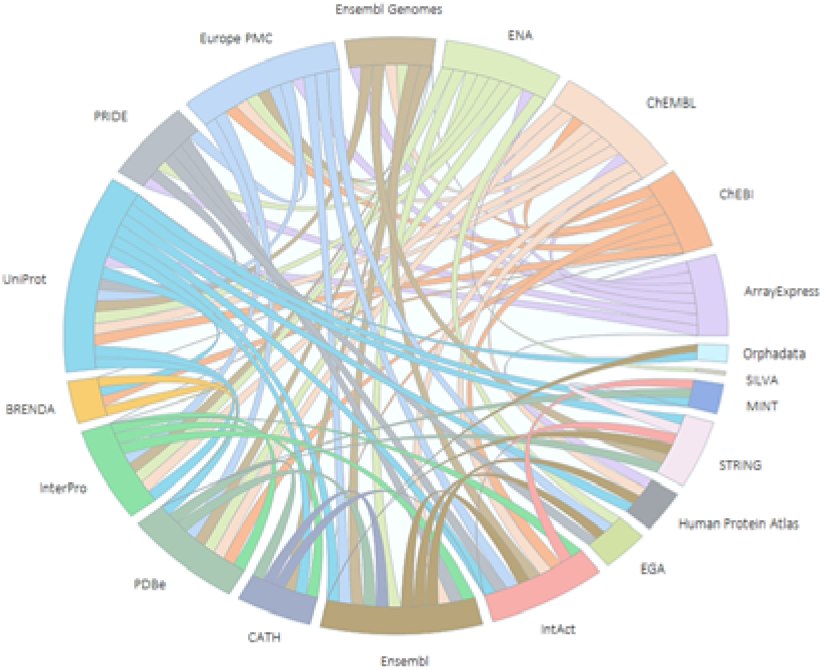
Core Data Resource interconnectivity.

Figure 5: *Heat map of Core Data Resource co-citation*: This is related to Indicator 2c “Usage in research as measured through citation in the literature” from Durinx *et al.*, 2016. The citations of CDRs were collected as for Figure 3. For each unique PMID across the three citation indicators, Cited-by counts were retrieved from Europe PMC. For each pair of resources, the number of common unique PMIDs were counted and displayed graphically as the log of the co-citation count for those two resources. Although co-citations do occur across the full set of CDRs, for legibility only the 12 CDRs that are most co-cited are displayed in Figure 5.

**Figure 5.**
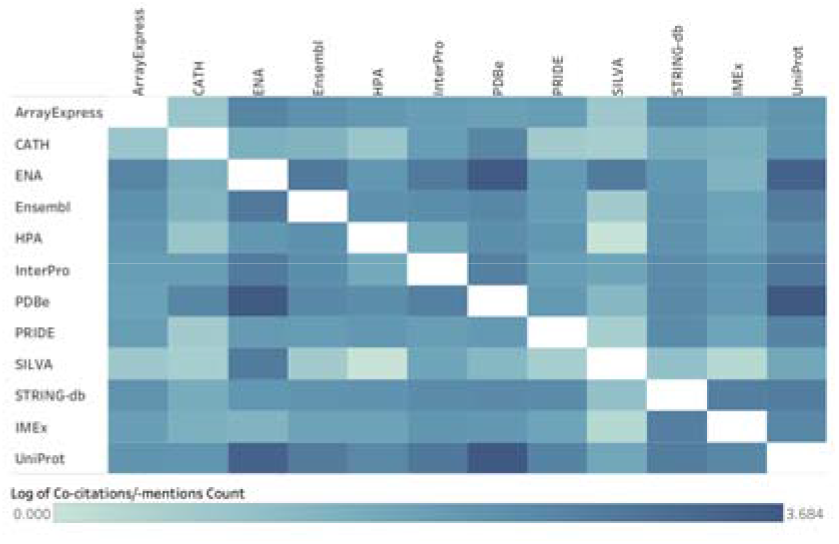
Heat map of the pairwise co-citation of the 12 ELIXIR Core Data Resources that are most frequently co-cited: ArrayExpress, CATH, ENA. Ensembl, HPA, InterPro, PDBe, PRIDE. SILVA, STRING-db, IMEx, and UniProt. The intensity of shading correlates with the frequency of co-citation.

Figure 6: *Horizon of assured funding*: This is related to Indicator 1d “Staff effort” from Durinx *et al.*, 2016. CDR managers were asked “As of January 2019, for how many Full Time Employees (FTEs) do you have committed funding, on 1 January in the following years?” Data were requested for 2019 to 2024. The figures reported do not imply that the baseline (January 2019) count reflects optimal staffing; resources may have been sub-optimally funded at the time of the survey. Nor do the figures imply that the resources anticipate that support will necessarily decline as shown — efforts to secure future funding are foremost in the minds of the resource managers, and ongoing. The survey question was intentionally specific, aiming to capture the assured security of staff funding for the infrastructure, projected forwards.

**Figure 6.**
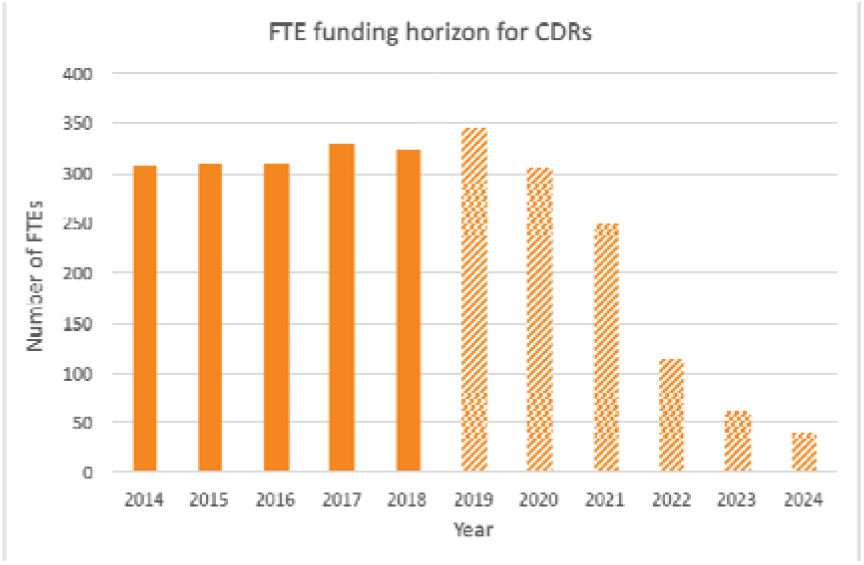
Horizon of assured funding: number of Full Time Equivalent positions at the CDRs from 2014-2018 (solid columns) and number of FTEs for which funding is assured 2019-2024 (striped columns), by year.

## 3 Results

Here, we describe the Core Data Resource infrastructure in terms of the scale of data housed, usage, and staffing, followed by a consideration of impact as reflected by citation data, and interdependencies and synergies in their inter-relationships. We reflect upon the role the CDRs play with respect to Open and FAIR data initiatives, and close with an examination of the security, in the long term, of the funding support on which they depend.

### 3.1 Scale of the Core Data Resources

Figure 1 shows the cumulative number of data entries across the Core Data Resources, including all deposited, curated and computed records. The total number of data entries almost quadrupled, from 967 million to 3,809 million, between 2014 and 2018.

The number of unique IP addresses accessing the data resources increased by 50 percent during the same period. As noted in the Methods, IP address figures are proxies for the number of individuals who use the CDRs. However, even with very conservative modelling (see discussion in https://beagrie.com/static/resource/EBI-impact-report.pdf) the number of scientists using the CDRs per month, given almost three million unique IP addresses, is in the hundreds of thousands. Additionally, we are confident that the increase in unique IP addresses is an indicator of real growth in users: this figure increased by more than 50 percent from 2014 to 2018.

Cumulative number of data entries in all Core Data Resources, plotted in conjunction with usage (as measured via the number of unique IP addresses accessing the CDRs per month), and the number of staff at the CDRs (as measured by Full Time Equivalents), per year.

How many people maintain, curate and serve these data to all these users? The number of FTEs employed in the Core Data Resources grew from 308 to 325, or just five percent, over the observed five-year period (Figure 1). Staff numbers are thus growing only slowly despite substantial increases in usage of the Core Data Resources and in their size as measured by the number of records and bytes (their “storage footprint”). This reflects the scalability of the technical solutions that have been adopted, the highly skilled workforce, and the value for money these resources offer. For each FTE employed, requests from almost 10,000 unique IP addresses per month are recorded.

Science evolves continually; developing data services such as metadata schemas, ontologies and user interfaces to support those evolving needs, while also maintaining backward compatibility to older data, is a distinctly human effort. Retaining and finding talented and knowledgeable staff to maintain the scientific relevance of CDRs and support their continued growth in usage requires continual investment.

### 3.2 Core Data Resource Citations in the Scientific Literature

Citation in the scientific literature is an established indicator of the value and significance of scientific resources including databases (Bousfield *et al.*, 2016) and can be assessed via text mining methods (Duck *et al.*, 2016, Kafkas *et al.*, 2013) as well as by traditional citation counts. We investigated the impact of Core Data Resources by mining the full text open access publications available in Europe PMC for mentions of Core Data Resources by their name and by their specific data entry identifiers, with open citations of Key Articles describing each specific resource being also included in the analysis. Figure 2 shows the growth in the number of publications in Europe PMC on the basis of these three citation indicators.

Given the total of 51,434 name or data identifier mention citations in 2017, a year in which around 305,000 open access articles were published, 17 percent of the open access articles in Europe PMC refer to a Core Data Resource by mentioning the resource name or an entry identifier. This is a significant proportion. As shown in Figure 2 (and Table S3 in Supplementary Data), the combined citation indicators for the Core Data Resources grew by 33 per cent, from 43,261 to 57,617, over the five-year period analysed.

Left axis: Sum of the number of mentions of the names of the resources (16 CDRs) and the resource entry identifiers (12 CDRs), per year, in the open access literature. Right axis: citations of pre-identified Key Articles describing the respective resources (18 CDRs). Note that citation data for 2018 are not depicted, as these were not available at the time the analysis for this figure was carried out.

Having established that the Core Data Resources are widely cited in the literature we assessed the scientific fields of the citing journals, by mapping the journals in which citations appear to Categories of scientific fields. As shown in Figure 3, the impact of the Core Data Resources beyond the immediate basic research domain from which they originated is clearly evident. The Core Data Resources are used more widely than within bioinformatics and molecular biology, ranging from primary research into applied and health sciences, food security and the environment.

### 3.3 Integration, Dependency and Ecosystem

The Core Data Resources exhibit high connectivity and interdependencies, reflecting the biological relationships between the different data types. The use of persistent identifiers is the primary method of cross-referencing between different resources, alongside the use of standard shared vocabularies such as the Gene Ontology (The Gene Ontology Consortium 2019). For example, UniProt protein sequences are translated from ENA sequences and Ensembl, and linked to corresponding PDBe structures. Records for compounds in ChEMBL link to IntAct interactions in which they are involved. The InterPro consortium builds on UniProt sequences to generate protein family signatures, which in turn are used to annotate uncharacterised UniProt sequence data. Most resources link to publications (Europe PMC) for biological context, which in turn cite identifiers to link back to the data. Figure 4 shows a representation of the interconnectivity between the CDRs. As new Core Data Resources are identified, it is expected that they will contribute to and extend this ecosystem.

While the CDRs support each other with the interconnections illustrated in Figure 4, they also interact with multiple resources outside this set. For example, ChEBI is used in UniProt enzyme annotations in the form of Rhea chemical reactions (https://www.rhea-db.org/), and UniProt enzymes are annotated using the IUBMB enzyme classification (https://iubmb.org/) as represented by the ENZYME database (https://enzyme.expasy.org/). While SILVA links to the ENA Core Data Resource, it also cross-references to RNACentral (https://rnacentral.org/), and the prokaryotic standard name resource LPSN (http://www.bacterio.net/) among others.

Between them, the Core Data Resources link to more than 350 external resources in more than 630 outward links in total, all listed in Table S6 in the Supplementary Data and Supporting Material; a deeper analysis of this data ecosystem in terms of scale, scope and interdependencies would be an interesting follow-on study. The diversity of this wider set of resources illustrates the foundational role of the Core Data Resources in the global bioinformatics landscape. Worldwide, the life science data resource ecosystem is an interlinked network, and the Core Data Resources are important nodes in that they integrate and make findable the data from hundreds of other resources, many of which are smaller, or domain-specific. In this way the Core Data Resources enhance the value of the other resources to which they are linked by multiplying re-use of their data.

The Core Data Resources are placed on the circumference of the circle, with each resource represented by an arc proportional to the total number of types of data directly exchanged between the two resources. The width of each internal arc, which transects the circle and connects two different resources, is proportional to the number of links between the two resources at the ends of the arc.

Another way to represent the integrated nature of Core Data Resources is to analyse the co-citation of different data resources in full text publications. That is, to count the number of times two or more resources (name or entry identifiers) are cited in the same publication. Figure 5 depicts the co-citation distribution for the 12 Core Data Resources that show the most co-citation. Notable co-citation hotspots include UniProt, PDBe and ENA, attesting to their frequent use in conjunction with each other and with other Core Data Resources.

### 3.4 Open Data and FAIR Data Leadership

Wide usage of data resources depends critically on the legal right to access and reuse data. This dependency is reflected in Indicator 4b “Open science” in the Core Data Resource selection process (Durinx *et al.*, 2016). All ELIXIR Core Data Resources are open access and enable the reuse and remixing of data, with either Terms of Use statements (12 of the resources) or specific licences (seven of the resources) that allow reuse. The Creative Commons licenses CC0, CC-BY or CC-BY-SA all conform to the Open Definition (http://opendefinition.org/licenses/), and serve for this purpose. Notably, as part of the process of application for Core Data Resource status, six resources amended their licences to being more permissive in order to fulfill the Open Science Indicator criterion.

ELIXIR supports European life scientists in making their data Findable, Accessible, Interoperable and Reusable (Blomberg and the ELIXIR Consortium, 2017). The Core Data Resources exemplify FAIR data across all FAIR principles (Wilkinson *et al.*, 2016). Durinx *et al.*, (2016) includes a Table within Box 1 that maps the Core Data Resource Indicators to FAIR Principles. For example, the “open data” characteristic of the Core Data Resources corresponds to the FAIR Principle “Reuseable”. Furthermore, Core Data Resources use persistent identifiers (Findable), standard vocabularies, and ontologies (Interoperable) as the norm in their metadata (included in the “Quality of service” Indicator 3a and 3d category). Data exchange is enabled via standard protocols such as HTTPS (websites and APIs) and FTP (“Quality of service” Indicator 3f, and FAIR principle “Accessible”). The Core Data Resources provide user support and customer service via helpdesks, user feedback mechanisms, and outreach and training activities (“Quality of service” Indicator 3g), to facilitate all aspects of the FAIR principles.

Wilkinson *et al.*, (2019) note that

> *The FAIR Principles are aspirational in that they do not strictly define how to achieve a state of ‘FAIRness’, rather they describe a continuum of features, attributes, and behaviors that move a digital resource closer to that goal. Despite their rapid community uptake, the question of how the FAIR Principles should be implemented has been prone to diverse interpretation. Some resource providers claim to be ‘already FAIR’ or ‘to enable FAIRness’ - statements that currently cannot be objectively evaluated.*

Although by definition the Core Data Resources are well-established and thus pre-date the advent of the FAIR Principles, several of the CDRs contribute to initiatives exploring the practicalities of FAIR Implementation. Examples include the Bioschemas community initiative (https://bioschemas.org/), which aims to improve the Findability of data in the life sciences by encouraging the use of Schema.org markup in data resource websites, and an ELIXIR Implementation Study “FAIRness of the current ELIXIR Core resources: Application (and test) of newly available FAIR metrics, and identification of steps to increase interoperability” (https://elixir-europe.org/platforms/data/fairness-core-resources).

### 3.5 Funding Horizon

ELIXIR Core Data Resources are the repository of record for a number of data types. Funders, journals, and submitters treat the Core Data Resources as stably funded infrastructure, but funding is in fact not assured past a very short horizon for many resources.

To assess the magnitude of this problem we asked managers of each Core Data Resource to report the funding for their staff that is currently confirmed. Figure 6 shows that based on data gathered early in 2019, the resources have assured funding for on average 88 percent of the staff within a one-year horizon to 2020, but on average only 33 percent of the staff over a three-year horizon to 2022. Just five of the 19 resources (26 percent) have the assurance that, one year from January 2019, they would have funds to support the same level of staffing as on that date.

These results show that beyond 2020 the assured funding levels are not sustained, implying a clear risk to the continued existence of this essential research infrastructure. The lack of assured long-term support for these mature and foundational resources demonstrates the fragility of data infrastructure upon which the research ecosystem depends and upon which funding agencies rely for storing research data generated with public monies.

It is unlikely, of course, that staffing for the infrastructure will actually collapse on the trajectory shown in Figure 6, as all Core Data Resources have demonstrated their capability to secure adequate funding for their operations, to date. However funding for much of the infrastructure is currently awarded on the basis of short-term grants or contracts whose terms are often suited more to research projects than to funding infrastructure. Consequently, resource managers spend a significant proportion of their time demonstrating the value of their resources to funders and preparing applications for funding renewal. It is entirely appropriate for funders to exercise mechanisms that continually assess the fit of the infrastructure with the scientific need, but Figure 6 suggests that the frequency of this assessment is faster than might be warranted for an infrastructure, which by definition must be established, of proven utility, and stable over time.

## 4 Discussion

During the past four decades, the massive growth of data in life sciences research, and the demonstration by researchers and funders that these data are more valuable if shared and re-used, have led to the creation of thousands of data resources to store, curate, and share these data (Imker, 2018; http://bigd.big.ac.cn/databasecommons/). Together, these data resources represent a new type of research infrastructure, which, unlike traditional “bricks and mortar” research infrastructures, is both virtual and distributed. The resources that make up this infrastructure are developed and maintained through the expertise of highly qualified personnel. The physical components of the infrastructure are these staff and the computational resources within which the data are stored and through which they are distributed to users.

The successful selection by ELIXIR of a set of Core Data Resources for Europe has shown that it is possible to develop a data-driven process to measure the impact of data resources and to use this process to identify a subset of those resources from within the larger data resource ecosystem that are most crucial to the larger infrastructure. The ELIXIR Core Data Resources define a cohort within the global life sciences infrastructure that funders and other stakeholders may use as a basis for structuring policies that support long-term sustainability, for both the Core Data Resources and the greater worldwide life sciences data infrastructure of which they are a part.

In addition to making the case for more sustainable funding support, the named Core Data Resources are models of good practice for managing data resources. They provide a focus for initiatives to integrate data and workflows from other, smaller data resources. Several of the Core Data Resources serve as the repository of record for archiving the data type they store: they are crucially important for the long term preservation of hard-won experimental data generated with public funding. The selection process itself provides a basis for selecting exemplars of good practice for other resource types, such as ELIXIR’s Recommended Interoperability Resources (https://www.elixir-europe.org/platforms/interoperability/rirs), as part of building the European research infrastructure across all components necessary for life sciences research.

The Core Data Resources identified by ELIXIR are, by definition, of fundamental importance to the life sciences research infrastructure in Europe and the rest of the world, and, for the first time here, this assertion is quantitatively demonstrated across the set of Core Data Resources. We have shown that these Core Data Resources are accessed by hundreds of thousands of users per month (Figure 1); they are explicitly cited in 17 percent of open access publications in Europe PMC (Figure 2); and they are used extensively across all fields in academic life sciences, medical sciences, and in various life sciences-related commercial activities (Figure 3). It is clear from our analysis that the value of the Core Data Resources infrastructure for the scientific research effort is continually increasing over time as archived data and the use of those data grows.

While the ELIXIR Core Data Resources are necessarily mature and stable, by virtue of the selection criteria that define them, Periodic Review of the CDR set is needed. This ensures that the standards applied in making the current selections are maintained, as is the relevance of the scope of the chosen resources to the life sciences, given that new technologies will be developed and new fields of research will emerge, in some cases superseding older technologies, and refocusing research priorities. In line with plans indicated in Durinx *et al.*, (2016), the first formal Periodic Review is due to take place in 2020.

### Risk to this critical infrastructure

The data infrastructure exemplified by the Core Data Resources has become essential to life sciences research worldwide, as well as in more applied settings such as healthcare, environmental science, biotechnology and food science, and operates in the commercial sector such as the pharmaceutical industry and many small-to-medium-sized companies (https://f1000research.com/documents/7-590). In recognition of the underpinning nature of open data to both research and the science-driven economy, virtually all research funders, both public and charitable, now strongly recommend or require deposition of research data into open access data resources (European Research Council: https://erc.europa.eu/sites/default/files/document/file/ERC_info_document-Open_Research_Data_and_Data_Management_Plans.pdf; Science Europe: https://www.scienceeurope.org/wp-content/uploads/2018/01/SE_Guidance_Document_RDMPs.pdf). Leading scientific journals, addressing their concerns about research reproducibility, increasingly advocate and, in some cases, require deposition into open access data repositories of research data associated with the articles they publish (*Scientific Data*: https://www.nature.com/sdata/policies/data-policies; PLOS: https://journals.plos.org/plosone/s/data-availability). Consequently, the core resources in this global enterprise require correspondingly sustainable funding (Bourne *et al*., 2015; Anderson *et al.* 2017).

The need for increased sustainability of funding for the ELIXIR Core Data Resources is demonstrated in Figure 6. Support to date, as shown by funding for FTEs from 2014-2018, appears stable. These figures, though, mask the fragility of this funding, which is heavily reliant upon short-term funding for which there is no guarantee of continuation, as shown by the sharp decline in assured funding for FTEs from 2020. Partly because they have been so successful in securing their funding to date, there is an under-appreciation of the precariousness of the funding landscape for such data resources, as evidenced by common assumptions that funding is likely to continue (Siepel, 2019). This contrasts with explicit acknowledgement that the landscape is changing even for well-established data resources, such as those supported by the NIH’s National Human Genome Research Institute (Kaiser, 2016) and neglects the fact that emerging research directions need to be incorporated, ensuring best service to the life science community. Consequently, solutions are needed to ensure adequate provision of foundational bioinformatics data resources in the long term.

### Worldwide data ecosystem

The European resources from which ELIXIR Core Data Resources are selected represent only a fraction of life sciences data resources worldwide. The rest of the world also develops and hosts data resources, and many of these are as important to the global life sciences data ecosystem as are the ELIXIR Core Data Resources. Indeed, several of the ELIXIR Core Data Resources are already members of international consortia, with ENA (INSDC; http://www.insdc.org/), PDBe (wwPDB; https://www.wwpdb.org/), and UniProt (https://www.uniprot.org/) being three prominent examples. Many of the global resources are also at risk from short-term and unstable funding cycles. The ELIXIR Core Data Resource selection process provides a model for identification of other crucial resources worldwide that will allow funders to more efficiently support the worldwide life sciences data resource ecosystem. The nascent Global Biodata Coalition (Anderson 2017; Anderson *et al.* 2017), supported by national and charitable funders globally, will use this process as a stepping stone towards a worldwide effort to identify and secure long-term funding for crucial data resources.

## Acknowledgements

The authors thank all contributors to the Core Data Resources, and John Hancock as well as the three reviewers for valuable comments on the manuscript.

## References

Anderson W.P., Global Life Science Data Resources Working Group. (2017) Data management: A global coalition to Sustain core data. Nature, 543, 179.

Athar A. et al. (2019) ArrayExpress update - from bulk to single-cell expression data. Nucleic Acids Res; 47(D1), D711–D715.

Berman H.M. (2008) The Protein Data Bank: a historical perspective. Acta Crystallogr A, 64, 88–95.

Blomberg N and ELIXIR Consortium. ELIXIR position paper on FAIR data management in the life sciences [version 1; not peer reviewed]. F1000Research 2017, 6(ELIXIR):1857

Bourne P.E. et al. (2015) Perspective: Sustaining the big-data ecosystem. Nature, 527, S16–17.

Bousfield D. et al. (2016) Patterns of database citation in articles and patents indicate long-term scientific and industry value of biological data resources. F1000Res, 5(ELIXIR), 160.

Cunningham F. et al. (2019) Ensembl 2019. Nucleic Acids Res, 47(D1), D745–D751.

Duck G., et al. (2016) A Survey of Bioinformatics Database and Software Usage through Mining the Literature. PLoS One. 11:e0157989

Durinx C. et al. (2016) Identifying ELIXIR Core Data Resources. F1000Res, 5(ELIXIR), 2422.

Gabella C. et al. (2017) Funding knowledgebases: Towards a sustainable funding model for the UniProt use case. F1000Res, 6(ELIXIR), 2051.

Glöckner F.O, et al. (2017) 25 years of serving the community with ribosomal RNA gene reference databases and tools. J Biotechnol, 261, 169–176.

Harrison P.W. et al. (2019) The European Nucleotide Archive in 2018. Nucleic Acids Res, 47(D1), D84–D88.

Hastings J. et al. (2016) ChEBI in 2016: Improved services and an expanding collection of metabolites. Nucleic Acids Res, 44(D1), D1214–1219.

Imker, H.J. (2018) 25 Years of Molecular Biology Databases: A Study of Proliferation, Impact, and Maintenance. Frontiers in Research Metrics and Analytics 8, 3.

Jeske L. et al. (2019) BRENDA in 2019: a European ELIXIR core data resource. Nucleic Acids Res, 47(D1), D542–D549.

Kafkas Ş., et al. (2013) Database citation in full text biomedical articles. PLoS One, 8:e63184

Kaiser J. (2016) Funding for key data resources in jeopardy.Science 351(6268):14.

Kersey P.J. et al. (2018) Ensembl Genomes 2018: an integrated omics infrastructure for non-vertebrate species. Nucleic Acids Res, 46(D1), D802–D808.

Lappalainen I. et al. (2015) The European Genome-phenome Archive of human data consented for biomedical research.Nat Genet, 47(7), 692–695.

Levchenko M. et al. (2018) Europe PMC in 2017. Nucleic Acids Res, 46(D1), D1254–D1260.

Mendez D. et al. (2019) ChEMBL: towards direct deposition of bioassay data. Nucleic Acids Res, 47(D1), D930–D940.

Metwally A. and Paduano M. (2011) Estimating the number of users behind ip addresses for combating abusive traffic. Proceedings of the 17th ACM SIGKDD international conference on Knowledge discovery and data mining 249–257.

Metwally A. et al. (2014) Large-Scale Network Traffic Analysis for Estimating the Size of IP Addresses and Detecting Traffic Anomalies. CRC Press 2014, Chapter 14, 435–462.

Mir S. et al. (2018) PDBe: towards reusable data delivery infrastructure at protein data bank in Europe. Nucleic Acids Res, 46(D1), D486–D492.

Mitchell A.L. et al. (2019) InterPro in 2019: improving coverage, classification and access to protein sequence annotations. Nucleic Acids Res, 47(D1), D351–D360.

Orchard S. et al. (2012) Protein interaction data curation: the International Molecular Exchange (IMEx) consortium. Nat Methods, 9 (4), 345–350.

Perez-Riverol Y. et al. (2019) The PRIDE database and related tools and resources in 2019: improving support for quantification data. Nucleic Acids Res, 47(D1), D442–D450.

Rath, A. et al. (2012) Representation of rare diseases in health information systems: the Orphanet approach to serve a wide range of end users. Hum Mutat, 33 (5), 803–808.

Siepel, A. (2019) Challenges in funding and developing genomic software: roots and remedies. Genome Biology 20:147.

Sillitoe I. et al. (2019) CATH: expanding the horizons of structure-based functional annotations for genome sequences. Nucleic Acids Res, 47(D1), D280–D284.

Szklarczyk D. et al. (2019) STRING v11: protein-protein association networks with increased coverage, supporting functional discovery in genome-wide experimental datasets. Nucleic Acids Res, 47(D1), D607–D613.

The Gene Ontology Consortium. (2019) The Gene Ontology Resource: 20 years and still GOing strong. Nucleic Acids Res, 47(D1), D330–D338.

Uhlén M. et al. (2015) Tissue-based map of the human proteome. Science, 347(6220), 1260419.

UniProt Consortium. (2019) UniProt: a worldwide hub of protein knowledge. Nucleic Acids Res, 47(D1), D506–D515.

Wilkinson M.D. et al. (2016) The FAIR Guiding Principles for scientific data management and stewardship. Sci Data, 3, 160018.

Wilkinson M.D. et al. (2019) Evaluating FAIR maturity through a scalable, automated, community-governed framework. Sci Data, 6, 174.

