## Supplementary Data for "The ELIXIR Core Data Resources: fundamental infrastructure for the life sciences"

The “Supporting Material” referred to within this Supplementary Data can be found in Supporting.Material.CDR.infrastructure.12082019.xlsx, DOI: 10.5281/zenodo.3468133 (https://zenodo.org/record/3468133).

**Figure 1. Scale of the Core Data Resources**

**Table S1. Data from which Figure 1 is derived:**

| **Year** | **2014** | **2015** | **2016** | **2017** | **2018** |
| --- | --- | --- | --- | --- | --- |
| **Data entries** | 967,333,131 | 1,738,151,018 | 1,853,898,281 | 2,838,070,639 | 3,808,683,537 |
| **Monthly user/IP addresses** | 2,107,904 | 2,408,037 | 2,504,140 | 2,960,632 | 3,209,646 |
| **FTEs** | 308.4 | 311.2 | 309.9 | 330.2 | 325.2 |

Figure 1 includes data from the following Core Data Resources:

ArrayExpress, BRENDA, CATH, ChEBI, ChEMBL, EGA, ENA, Ensembl, Ensembl Genomes, EuropePMC, HPA, IntAct /MINT , InterPro, Orphadata, PDBe, PRIDE, SILVA, STRING, UniProt

- Note that Ensembl’s compute infrastructure physically relocated in 2016, so “Users/IP address” data are not available for that year. In this case, the 2015 numbers were rolled forward to 2016.
- Note that STRING made only minor releases in 2014, 2016 and 2018, in that the interactions are re-computed, but the number of “Data entries” remains unchanged. The major releases that change the number of “Data entries” happened in 2013, 2015 and 2017. So, for “Data entries”, the total reported for 2013 was rolled forward to 2014, the total for 2015 was rolled forward to 2016, and the total for 2017 was rolled forward to 2018.
- Note that SILVA did not release a major update in 2018, so for “Data Entries” the total for 2017 was rolled forward to 2018.

**Figure 2: Usage of Core Data Resources in research**

The following steps were taken:

1. API calls were run on *open access full text articles* in Europe PMC to identify articles that mention Core Data Resource by name or include specific data record accession numbers (see Venkatesan et al., 2017, “SciLite: a platform for displaying text-mined annotations as a means to link research articles with biological data” (https://doi.org/10.12688/wellcomeopenres.10210.2), and http://europepmc.org/developers).
   a. The identification is based on pattern matching for each CDR as described at <https://github.com/EuropePMC/EuropePMC-Identifier-Extractor/blob/master/automata/resources170731.mwt>

and

<https://github.com/EuropePMC/EuropePMC-Identifier-Extractor/blob/master/automata/acc181210.mwt>

b. The URL for each CDR API call is below. Each RESOURCE_NAME API call searches multiple strings representing variants of each CDR name, as listed in the links in point (a) above.

Resource Name mentions:

ArrayExpress - <http://europepmc.org/search?query=RESOURCE_NAME:arrayexpress>

CATH - <http://europepmc.org/search?query=RESOURCE_NAME:cath>

ChEBI - <http://europepmc.org/search?query=RESOURCE_NAME:chebi>

ChEMBL - <http://europepmc.org/search?query=RESOURCE_NAME:chembl>

EGA - <http://europepmc.org/search?query=RESOURCE_NAME:ega>

ENA - <http://europepmc.org/search?query=RESOURCE_NAME:ena>

Ensembl - <http://europepmc.org/search?query=RESOURCE_NAME:ensembl>

Ensembl Genomes - <http://europepmc.org/search?query=RESOURCE_NAME:ensemblgenomes>

Europe PMC - <http://europepmc.org/search?query=RESOURCE_NAME:epmc>

HPA - <http://europepmc.org/search?query=RESOURCE_NAME:hpa>

IntAct - <http://europepmc.org/search?query=RESOURCE_NAME:intact>

MINT - <http://europepmc.org/search?query=RESOURCE_NAME:mint>

InterPro - <http://europepmc.org/search?query=RESOURCE_NAME:interpro>

PDBe - <http://europepmc.org/search?query=RESOURCE_NAME:pdb>

PRIDE - <http://europepmc.org/search?query=RESOURCE_NAME:pride>

STRING-db - <http://europepmc.org/search?query=RESOURCE_NAME:stringdb>

UniProt - <http://europepmc.org/search?query=RESOURCE_NAME:uniprot>

Accession Number mentions:

ArrayExpress - <http://europepmc.org/search?query=ACCESSION_TYPE:arrayexpress>

CATH - <http://europepmc.org/search?query=ACCESSION_TYPE:cath>

ChEBI - <http://europepmc.org/search?query=ACCESSION_TYPE:chebi>

ChEMBL - <http://europepmc.org/search?query=ACCESSION_TYPE:chembl>

EGA - <http://europepmc.org/search?query=ACCESSION_TYPE:ega>

ENA - <http://europepmc.org/search?query=ACCESSION_TYPE:ena>

Ensembl - <http://europepmc.org/search?query=ACCESSION_TYPE:ensembl>

HPA - <http://europepmc.org/search?query=ACCESSION_TYPE:hpa>

IntAct - <http://europepmc.org/search?query=ACCESSION_TYPE:intact>

MINT - <http://europepmc.org/search?query=ACCESSION_TYPE:mint>

InterPro - <http://europepmc.org/search?query=ACCESSION_TYPE:interpro>

PDBe - <http://europepmc.org/search?query=ACCESSION_TYPE:pdb>

PRIDE - <http://europepmc.org/search?query=ACCESSION_TYPE:pxd>

UniProt - <http://europepmc.org/search?query=ACCESSION_TYPE:uniprot>

c. The counts from the API calls were totalled, for each year between 2013 and 2017 inclusive (top panel, Table S3 below)

2. For each CDR selected Key Article, listed by PMID in Table S2 below, the following API call was made, to tally the number of citations for that PMID:
   [https://www.ebi.ac.uk/europepmc/webservices/rest/MED/**<pmid>**/citations?page=<**num>**&pageSize=**1000**&format=json](https://www.ebi.ac.uk/europepmc/webservices/rest/MED/25361974/citations?page=1&pageSize=1&format=json)

From this output (reported in the Fig2.cdr_citations_25Jan2019 tab in the Supporting Material [here](https://zenodo.org/record/2625247)) the number of citing articles was counted for each year between 2013 and 2017 inclusive (middle panel, Table S3 below)

3. The results from Steps 1 and 2 were aggregated (bottom panel, Table S3 below) and used to draw the Figure 2 graphic.

**Table S2. Key Article PMIDs.**

| **Database** | **PMIDs of Key Articles** |
| --- | --- |
| ArrayExpress | 12519949  14744115 15608260 16939801 19015125 21071405 23193272 25361974 |
| BRENDA | 11796225 17202167 18984617 21030441 21062828 23203881 25378310 27924025 |
| CATH | 17135200 18996897 19679085 19758469 20368142 21097779 25348408 26139634 26253692 27899584 28150234 |
| ChEBI | 17932057 19496059 19854951 23180789 26467479 |
| ChEMBL | 23657106 24214965 24635517 25883136 26201396 27899562 28602100 |
| EGA | 26111507 |
| ENA | 20972220 23203883 24214989 25404130 26615190 27899630 29140475 |
| Ensembl | 24316576 25352552 27141089 27268795 27337980 27899575 29155950 |
| Ensembl Genomes | 19884133 22067447 24163254 24217918 25432969 26578574 |
| Europe PMC | 23734176 25378340 25774284 25789152 28948232 29161421 |
| Human Protein Atlas | 16127175 21139605 25613900 |
| InterPro | 17202162 18940856 22096229 24451626 25428371 27899635 |
| PDBe | 28573592 29126160 29174494 29533231 29749603 |
| PRIDE | 16041671 16381953 19662629 19906717 23203882 27683222 |
| SILVA | 17947321 23193283 24293649 28648396 |
| STRING | 17098935 18940858 21045058 23203871 25352553 27924014 |
| The IMEx Consortium | 14681455 17135203 17145710 19850723 22096227 22121220 22453911 24234451 |
| UniProt | 21447597 22102590 24253303 25348405 27899622 29425356 |

**Table S3. Data from which Figure 2 is derived:**

|  | **Combined Resource Name and Accession Mention Counts*** | | | | |  |
| --- | --- | --- | --- | --- | --- | --- |
| **Year** | **2013** | **2014** | **2015** | **2016** | **2017** | **Grand Total** |
| **Total** | 40,653 | 42,872 | 46,712 | 48,424 | 51,434 | 230,095 |
| * For a single CDR, a PMID that mentions a resource name and its accession is counted twice. A PMID that mentions a resource name and an accession for two different CDRs is counted four times. | | | | | | |
|  | **CDR Key Article Citation Counts*** | | | | |  |
| **Year** | **2013** | **2014** | **2015** | **2016** | **2017** | **Grand Total** |
| **Total** | 2,608 | 3,223 | 4,561 | 5,411 | 6,183 | 21,986 |
| * A PMID that cites CDR Key Articles of two different CDRs is counted twice. | | | | | | |
|  | **Combined Resource Name, Accession Mention and CDR Key Article Citation Counts** | | | | |  |
| **Year** | **2013** | **2014** | **2015** | **2016** | **2017** | **Grand Total** |
| **Total** | 43,261 | 46,095 | 51,273 | 53,835 | 57,617 | 252,081 |

Figure 2 Resource Name/Accession mentions includes data from the following Core Data Resources: ArrayExpress, CATH, ChEBI, ChEMBL, EGA, ENA, Ensembl, Ensembl Genomes (name mentions only, not data accessions), EuropePMC (name mentions only, not data accessions), HPA, IntAct /MINT, InterPro, PDBe, PRIDE, STRING(name mentions only, not data accessions), UniProt. BRENDA, SILVA and Orphadata were not included in the initial list of Core Data Resources, and have not yet been folded into the “Resource Name Mentions” text mining pipeline.

Figure 2 Citation of Key Article counts uses data from the following Core Data Resources:

ArrayExpress, BRENDA, CATH, ChEBI, ChEMBL, EGA, ENA, Ensembl, Ensembl Genomes, Europe PMC, Human Protein Atlas, IntAct and MINT for The IMEx Consortium, InterPro, PDBe, PRIDE, SILVA, STRING, UniProt

**Figure 3. Categories of the scientific fields in which the 20 journals that most frequently cite the Core Data Resources are active.**

The following steps were taken, with data collections steps 1-3 being executed on 15th August 2018:

1. For each CDR selected Key Article, PMIDs listed in Table S2 above, the following API call was made to tally the number of citations for that PMID (e.g. [https://www.ebi.ac.uk/europepmc/webservices/rest/MED/**<pmid>**/citations?page=<**num>**&pageSize=**1000**&format=json](https://www.ebi.ac.uk/europepmc/webservices/rest/MED/25361974/citations?page=1&pageSize=1&format=json))
2. CDR resource name mention PMIDs were then collected via Europe PMC’s APIs using resource-specific search patterns:

**Table S4. Resource-specific search patterns.**

| **Core Data Resource** | **Search Pattern** |
| --- | --- |
| ArrayExpress | %22ArrayExpress%22 |
| ArrayExpress | %22Array Express%22 |
| BRENDA | %22BRENDA Tissue Ontology%22 |
| CATH | Protein Structure Classification CATH |
| ChEBI | %22ChEBI%22 |
| ChEMBL | %22ChEMBL%22 |
| EGA | European Genome-phenome Archive EGA |
| ENA | European Nucleotide Archive ENA |
| Ensembl | %22Ensembl%22 |
| Ensembl Genomes | %22Ensembl Genomes%22 |
| Ensembl Genomes | %22EnsemblGenomes%22 |
| Ensembl Genomes | %22Ensembl Metazoa%22 |
| Ensembl Genomes | %22EnsemblMetazoa%22 |
| Ensembl Genomes | %22Ensembl Plants%22 |
| Ensembl Genomes | %22EnsemblPlants%22 |
| Ensembl Genomes | %22Ensembl Protists%22 |
| Ensembl Genomes | %22EnsemblProtists%22 |
| Ensembl Genomes | %22Ensembl Fungi%22 |
| Ensembl Genomes | %22EnsemblFungi%22 |
| Ensembl Genomes | %22Ensembl Bacteria%22 |
| Ensembl Genomes | %22EnsemblBacteria%22 |
| Europe PMC | %22Europe PMC%22 |
| Europe PMC | %22EuropePMC%22 |
| Human Protein Atlas | %22Human Protein Atlas%22 |
| InterPro | %22InterPro%22 |
| PDBe | %22PDBe%22 |
| PDBe | %22Protein Data Bank in Europe%22 |
| PRIDE | proteomics identifications database PRIDE |
| SILVA | %22SILVA database%22 |
| STRING-db | %22STRING-db%22 |
| STRING-db | %22STRING db%22 |
| STRING-db | %22STRINGdb%22 |
| The IMEx Consortium | %22IMEx Consortium%22 |
| The IMEx Consortium | IntAct %22Molecular Interaction database%22 |
| The IMEx Consortium | %22Molecular INTeraction Database%22 |
| The IMEx Consortium | MINT %22Molecular Interaction database%22 |
| UniProt | %22UniProt%22 |
| UniProt | %22The Universal Protein Resource%22 |

3. Finally, data accession mention PMIDs were collected using Europe PMC’s APIs via [Europe PMC’s text-mined terms](ftp://ftp.ebi.ac.uk/pub/databases/pmc/TextMinedTerms/) (ftp://ftp.ebi.ac.uk/pub/databases/pmc/TextMinedTerms/) (N.B. Europe PMC does not collect accessions for Ensembl Genomes and STRINGbecause they re-use Ensembl accession numbers, and there is no way to distinguish them) (results recorded in the Fig3.5.Step3.mine.acc.15Aug2018 tab in the Supporting Material [here](https://zenodo.org/record/2625247))
4. For each unique PMID across the preceding 3 sets (from steps 1, 2 and 3) Journal Title, Publication Year and Cited-by count were retrieved from Europe PMC (results recorded in the Fig3.5.Step4.cdr_all tab in the Supporting Material [here](https://zenodo.org/record/2625247))
5. Each journal title was mapped to a set of Categories, based on freely available data retrieved from [Scimago Journal & Country Rank (https://www.scimagojr.com/journalrank.php)](https://www.scimagojr.com/journalrank.php) (recorded in the Fig3.Step5.scimagojr_2016 tab in the Supporting Material [here](https://zenodo.org/record/2625247))
4. [6. ](https://www.scimagojr.com/journalrank.php)Based on the CDR-citing PMIDs identified in steps 1, 2 and 3, and the Journal Titles associated with them, the Top 20 CDR-citing journals were identified:

Antimicrobial Agents and Chemotherapy
Applied and Environmental Microbiology
Biochemistry
Biophysical Journal
BMC Genomics
Emerging Infectious Diseases
Frontiers in Microbiology
Genetics
Genome Announcements
Infection and Immunity
Journal of Bacteriology
Journal of Clinical Microbiology
Journal of Virology
Molecular and Cellular Biology
Nature Communications
Nucleic Acids Research
PLOS ONE
PLOS Pathogens
Proceedings of the National Academy of Sciences of the United States of America
Scientific Reports

For these 20 Journals, associated Category and corresponding Citation Count were extracted, and used to generate Figure 3.

**Table S5. Data from which Figure 3 is derived:**

| **Category** | **Citation Count** |
| --- | --- |
| Biochem, Genetics, Mol Biol (misc) | 22,945 |
| Multidisciplinary | 22,261 |
| Medicine (misc) | 20,609 |
| Agri and Biol Sciences (misc) | 20,609 |
| Genetics | 18,654 |
| Molecular Biology | 15,203 |
| Microbiology | 14,709 |
| Immunology | 12,356 |
| Biotechnology | 10,802 |
| Virology | 9,290 |
| Microbiology (medical) | 8,507 |
| Infectious Diseases | 8,417 |
| Applied Microbiol and Biotech | 7,515 |
| Food Science | 7,515 |
| Ecology | 7,515 |
| Parasitology | 4,784 |
| Pharmacology (medical) | 3,563 |
| Pharmacology | 3,563 |
| Cell Biology | 2,819 |
| Biochemistry | 2,447 |
| Physics and Astronomy (misc) | 2,336 |
| Biophysics | 2,313 |
| Epidemiology | 1,788 |

Figure 3 includes data from the following Core Data Resources:

ArrayExpress, BRENDA, CATH, ChEBI, ChEMBL, EGA, ENA, Ensembl, Ensembl Genomes, EuropePMC, HPA, IntAct/MINT, InterPro, PDBe, PRIDE, SILVA, STRING, UniProt.

**Figure 4. Core Data Resource interconnectivity, based on the reported links out from each CDR.**

The following steps were taken:

1. Each Core Data Resource was queried for a list of other data resources with which data are exchanged directly using computational links, for example via API, including other CDRs and also data resources beyond the CDR set. The links are shown in Table S6, gathered in March 2019.
2. For Figure 4, the relationships between Core Data Resources were expressed in a chord diagram, with the arc width weighted according to the number of outgoing links for each CDR.

**Table S6. Data from which Figure 4 is derived:**

| **Core Data Resource** | **Links to Other CDRs** | **Links to Additional Data Resources*** |
| --- | --- | --- |
| ArrayExpress | ChEBI  ChEMBL  ENA  Ensembl Genomes  Europe PMC  Human Protein Atlas  PRIDE  UniProt | BioSamples (https://www.ebi.ac.uk/biosamples/)  BioStudies (https://www.ebi.ac.uk/biostudies/)  EBI Mouse Resources (https://www.infrafrontier.eu/; http://www.mousephenotype.org/)  EFO (https://www.ebi.ac.uk/efo/)  EVA (https://www.ebi.ac.uk/eva/)  Expression Atlas (https://www.ebi.ac.uk/gxa/home)  GenomeSpace (http://www.genomespace.org/)  GEO (https://www.ncbi.nlm.nih.gov/geo/)  Identifiers.org (https://identifiers.org)  IGSR (http://www.internationalgenome.org/data-portal/sample)  MetaboLights (https://www.ebi.ac.uk/metabolights/)  MGnify (https://www.ebi.ac.uk/metagenomics/)  OmicsDI (https://www.omicsdi.org/)  Open Targets (https://www.opentargets.org/)  VectorBase (https://www.vectorbase.org/)  WormBase (https://www.wormbase.org) |
| BRENDA | ChEBI  Interpro  PDBe  UniProt | ENZYME (https://enzyme.expasy.org/)  ExplorEnz (https://www.enzyme-database.org/)  GENOME (https://www.ncbi.nlm.nih.gov/genome)  IUBMB enzyme nomenclature (https://iubmb.org/biochemical-nomenclature/)  KEGG (https://www.genome.jp/kegg/)  MetaCyc (https://metacyc.org/)  NCBI NUCLEOTIDE (https://www.ncbi.nlm.nih.gov/nucleotide/)  NCBI PROTEIN (https://www.ncbi.nlm.nih.gov/protein)  NCBI TAXONOMY (https://www.ncbi.nlm.nih.gov/guide/taxonomy/)  OMIM (https://www.omim.org/)  PubMed (https://www.ncbi.nlm.nih.gov/pubmed/)  SABIO-RK (http://sabiork.h-its.org/) |
| CATH | Ensembl  Ensembl Genomes  IntAct  InterPro  PDBe  UniProt | DrugBank (https://www.drugbank.ca/)  FunTree (http://www.funtree.info/FunTree/)  IntEnz (https://www.ebi.ac.uk/intenz/)  PDBsum (http://www.ebi.ac.uk/thornton-srv/databases/cgi-bin/pdbsum/)  Pfam (https://pfam.xfam.org/)  QuickGO (https://www.ebi.ac.uk/QuickGO/) |
| ChEBI | ChEMBL  BRENDA  Ensembl  Ensembl Genomes  Europe PMC  IntAct  InterPro  UniProt | Alan Wood's Pesticides (http://www.alanwood.net/pesticides/)  BioModels (http://www.ebi.ac.uk/biomodels/)  BioSamples (https://www.ebi.ac.uk/biosamples/)  BioStudies (https://www.ebi.ac.uk/biostudies/)  ChemIDplus (https://chem.nlm.nih.gov/chemidplus/)  Chemspider (http://www.chemspider.com/)  Complex Portal (https://www.ebi.ac.uk/complexportal/home)  DrugBank (https://www.drugbank.ca/)  DrugCentral (http://drugcentral.org/)  EBI Mouse Resources (https://www.infrafrontier.eu/; http://www.mousephenotype.org/)  ECMDB (http://ecmdb.ca/)  Enzyme Portal (https://www.ebi.ac.uk/enzymeportal/)  Expression Atlas (https://www.ebi.ac.uk/gxa/home)  FooDB (http://foodb.ca/)  GlyTouCan (https://glytoucan.org/)  Gmelin (http://202.127.145.151/siocl/cdbank/WebHelp/gmelin/gmehtml/pageintr.htm)  HMDB (http://www.hmdb.ca)  IntEnz (https://www.ebi.ac.uk/intenz/)  IPNI (https://www.ipni.org/)  KEGG (https://www.genome.jp/kegg/)  KNApSAcK (http://kanaya.naist.jp/KNApSAcK/)  LINCS (https://systemsbiology.columbia.edu/lincs)  LipidMaps (https://www.lipidmaps.org/)  MetaboLights (https://www.ebi.ac.uk/metabolights/)  MetaCyc (https://metacyc.org/)  MolBase (https://www.molbase.com/)  NCBI (https://www.ncbi.nlm.nih.gov/)  NIST Chemistry WebBook (https://webbook.nist.gov/chemistry/)  OLS (https://www.ebi.ac.uk/ols/index)  OmicsDI (https://www.omicsdi.org/)  Open Targets (https://www.opentargets.org/)  Patent (https://www.ebi.ac.uk/patentdata/proteins)  PDB (https://www.wwpdb.org/)  PubChem (https://pubchem.ncbi.nlm.nih.gov/)  Reactome (https://reactome.org)  Reaxys (https://www.elsevier.com/solutions/reaxys)  RESID (https://proteininformationresource.org/resid/)  SMID (http://www.smid-db.org/)  UM-BBD (https://www.hsls.pitt.edu/obrc/index.php?page=URL1100188151)  WebElements (https://www.webelements.com/)  Wikipedia (https://en.wikipedia.org/) |
| ChEMBL | ArrayExpress  BRENDA  ChEBI  Ensembl  Ensembl Genomes  EuropePMC  Human Protein Atlas  IntAct  InterPro  PDBe  UniProt | ACToR (https://actor.epa.gov/actor/home.xhtml)  BindingDB (https://www.bindingdb.org/bind/index.jsp)  BioModels (http://www.ebi.ac.uk/biomodels/)  BioStudies (https://www.ebi.ac.uk/biostudies/)  CanSAR (http://cansar.icr.ac.uk)  Carotenoid Database (http://carotenoiddb.jp/)  ClinicalTrials.gov (https://clinicaltrials.gov/)  Complex Portal (https://www.ebi.ac.uk/complexportal/home)  CREDO (http://marid.bioc.cam.ac.uk/credo)  DailyMed (https://dailymed.nlm.nih.gov/dailymed/)  DrugBank (https://www.drugbank.ca/)  DrugCentral (http://drugcentral.org/)  EBI Mouse Resources (https://www.infrafrontier.eu/; http://www.mousephenotype.org/)  EFO (https://www.ebi.ac.uk/efo/)  eMolecules (https://www.emolecules.com/)  Enzyme Portal (https://www.ebi.ac.uk/enzymeportal/)  EPA CompTox Dashboard (https://comptox.epa.gov/)  Expression Atlas (https://www.ebi.ac.uk/gxa/home)  FDA/USP SRS (https://fdasis.nlm.nih.gov/srs/)  Guide to Pharmacology (http://www.guidetopharmacology.org/)  GWAS Catalog (https://www.ebi.ac.uk/gwas/)  HPA (https://www.proteinatlas.org/)  Human Metabolome Database (http://www.hmdb.ca/)  Identifiers.org (https://identifiers.org)  KEGG (https://www.genome.jp/kegg/)  LINCS (https://systemsbiology.columbia.edu/lincs)  LipidMaps (https://www.lipidmaps.org/)  Mcule (https://mcule.com/)  MetaboLights (https://www.ebi.ac.uk/metabolights/)  MICAD (https://www.micad.co.uk/)  MolPort (https://www.molport.com/shop/index)  NIH Clinical Collection (http://nihsmr.evotec.com/evotec/)  Nikkaji (https://jglobal.jst.go.jp/en/)  NMRShiftDB (https://nmrshiftdb.nmr.uni-koeln.de/)  OLS (https://www.ebi.ac.uk/ols/index)  OmicsDI (https://www.omicsdi.org/)  Open Targets (https://www.opentargets.org/)  Pfam (https://pfam.xfam.org/)  PharmGKB (https://www.pharmgkb.org/)  Pharos (https://www.pharosproject.net/)  PubChem (https://pubchem.ncbi.nlm.nih.gov/)  Reactome (https://reactome.org)  Recon (https://reconchemicals.com/)  Rhea (https://www.rhea-db.org/)  Selleck (https://www.selleckchem.com/screening/fda-approved-drug-library.html)  TIMBAL (http://mordred.bioc.cam.ac.uk/timbal/)  VectorBase (https://www.vectorbase.org/)  Wikipedia (https://en.wikipedia.org/)  WormBase (https://www.wormbase.org)  ZINC (http://zinc15.docking.org/) |
| EGA | ENA  Ensembl  Europe PMC  PRIDE | BioSamples (https://www.ebi.ac.uk/biosamples/)  BioStudies (https://www.ebi.ac.uk/biostudies/)  dbGaP (https://www.ncbi.nlm.nih.gov/gap)  EBI Mouse Resources (https://www.infrafrontier.eu/; http://www.mousephenotype.org/)  EFO (https://www.ebi.ac.uk/efo/)  Expression Atlas (https://www.ebi.ac.uk/gxa/home)  ICGC (https://icgc.org/)  Identifiers.org (https://identifiers.org)  OLS (https://www.ebi.ac.uk/ols/index)  OmicsDI (https://www.omicsdi.org/)  RD-Connect (https://rd-connect.eu/)  Type 2 Diabetes Knowledge Portal (https://www.broadinstitute.org/diabetes/type-2-diabetes-knowledge-portal)  UK Biobank (https://mrc.ukri.org/research/facilities-and-resources-for-researchers/biobank/) |
| ENA | ArrayExpress  EGA  Ensembl  Ensembl Genomes  Europe PMC  InterPro  PDBe  PRIDE  SILVA  UniProt | BCCM/LMBP (http://bccm.belspo.be/about-us/bccm-lmbp)  BioModels (http://www.ebi.ac.uk/biomodels/)  BioSamples (https://www.ebi.ac.uk/biosamples/)  BioStudies (https://www.ebi.ac.uk/biostudies/)  CABRI (http://www.cabri.org/CABRI/srs-doc/cabi_fil.info.html)  CCAP (https://www.ccap.ac.uk/)  CNSA (https://db.cngb.org/cnsa/)  COMPARE-RefGenome (https://www.compare-europe.eu/library/reference-genomes)  DDBJ (https://www.ddbj.nig.ac.jp/)  dictyBase (http://dictybase.org/)  The Earlham Institute (http://www.earlham.ac.uk/services)  EBI Mouse Resources (https://www.infrafrontier.eu/; http://www.mousephenotype.org/)  EMBL Australian Bioinformatics Resource (https://www.embl-abr.org.au/)  EPD (https://epd.epfl.ch//index.php)  EVA (https://www.ebi.ac.uk/eva/)  Expression Atlas (https://www.ebi.ac.uk/gxa/home)  FlyBase (https://flybase.org/)  GeneDB (http://www.genedb.org/)  GFBIO (https://www.gfbio.org/)  GOA (https://www.ebi.ac.uk/GOA)  GrainGenes (https://wheat.pw.usda.gov/GG3/)  gtRNAdb (http://gtrnadb.ucsc.edu/)  GWAS Catalog (https://www.ebi.ac.uk/gwas/)  H-InvDB (http://www.h-invitational.jp/)  HGNC (https://www.genenames.org/)  Human Cell Atlas (https://www.humancellatlas.org/)  Identifiers.org (https://identifiers.org)  IGSR (http://www.internationalgenome.org/data-portal/sample)  IMGT/LIGM (http://www.imgt.org/ligmdb/documentation)  IntEnz (https://www.ebi.ac.uk/intenz/)  ISHAM-ITS (http://its.mycologylab.org/)  lncRNAdb (http://www.lncrnadb.org/)  MarCat (https://mmp.sfb.uit.no/databases/marcat/#/)  MarDB (https://mmp.sfb.uit.no/databases/mardb/)  MarRef (https://mmp.sfb.uit.no/databases/marref/)  MG-RAST (https://www.mg-rast.org/)  MGI (http://www.informatics.jax.org/)  MGnify (https://www.ebi.ac.uk/metagenomics/)  mirBase (http://www.mirbase.org/)  MPI Toolkit (https://toolkit.tuebingen.mpg.de/#/)  NCBI (https://www.ncbi.nlm.nih.gov/)  OLS (https://www.ebi.ac.uk/ols/index)  OmicsDI (https://www.omicsdi.org/)  PANGAEA (https://www.pangaea.de/)  PDB (https://www.wwpdb.org/)  PLncDB (https://omictools.com/plncdb-tool)  PomBase (https://www.pombase.org/)  PR2 (https://github.com/pr2database/pr2database)  Rfam (http://rfam.xfam.org/)  RNAcentral (https://rnacentral.org)  SequenceAnalysis.co.uk (http://sequenceanalysis.co.uk/)  SGD (https://www.yeastgenome.org/)  snOPY (http://snoopy.med.miyazaki-u.ac.jp/)  SRPDB (https://rth.dk/resources/rnp/SRPDB/)  SubtiList (http://genolist.pasteur.fr/SubtiList/)  TAIR (https://www.arabidopsis.org/)  tmRNA Website (https://bioinformatics.sandia.gov/tmrna/)  Unite (https://unite.ut.ee/)  VBASE2 (http://www.vbase2.org/)  VectorBase (https://www.vectorbase.org/)  VEGA (https://www.sanger.ac.uk/science/tools/vega-genome-browser)  WormBase (https://www.wormbase.org)  WoRMS (http://www.marinespecies.org/)  ZFIN (https://zfin.org) |
| Ensembl | CATH  ChEMBL  EGA  ENA  Ensembl Genomes  Europe PMC  Human Protein Atlas  IntAct  InterPro  Orphadata  PDBe  PRIDE  STRING  UniProt | Aniseed (https://www.aniseed.cnrs.fr/)  APPRIS (http://appris.bioinfo.cnio.es/#/)  BioModels (http://www.ebi.ac.uk/biomodels/)  BioSamples (https://www.ebi.ac.uk/biosamples/)  dbGaP (https://www.ncbi.nlm.nih.gov/gap)  Diana TarBase (http://carolina.imis.athena-innovation.gr/diana_tools/web/index.php?r=tarbasev8%2Findex)  DPGP (http://dpgp.org/)  EBI Mouse Resources (https://www.infrafrontier.eu/; http://www.mousephenotype.org/)  EFO (https://www.ebi.ac.uk/efo/)  ESP (http://evs.gs.washington.edu/EVS/)  EVA (https://www.ebi.ac.uk/eva/)  Expression Atlas (https://www.ebi.ac.uk/gxa/home)  FANTOM5 (http://fantom.gsc.riken.jp/5/)  GEFOS (http://www.gefos.org/)  GIANT (http://giant.princeton.edu/)  GOA (https://www.ebi.ac.uk/GOA)  GTEx (https://gtexportal.org/home/)  GWAS Catalog (https://www.ebi.ac.uk/gwas/)  HGNC (https://www.genenames.org/)  HPA (https://www.proteinatlas.org/)  Identifiers.org (https://identifiers.org)  IGSR (http://www.internationalgenome.org/data-portal/sample)  IHEC (http://ihec-epigenomes.org/)  IMPC (http://www.mousephenotype.org/)  JASPAR (http://jaspar.genereg.net/)  MAGIC (https://www.magicinvestigators.org/)  MGI (http://www.informatics.jax.org/)  mirBase (http://www.mirbase.org/)  NCBI (https://www.ncbi.nlm.nih.gov/)  OLS (https://www.ebi.ac.uk/ols/index)  OMIA (https://omia.org/home/)  OMIM (https://www.omim.org/)  Open Targets (https://www.opentargets.org/)  Orphanet (https://www.orpha.net/)  Pfam (https://pfam.xfam.org/)  Reactome (https://reactome.org)  RefSeq (https://www.ncbi.nlm.nih.gov/refseq/)  Rfam (http://rfam.xfam.org/)  RGD (https://rgd.mcw.edu/)  RNAcentral (https://rnacentral.org)  Sanger CRISPR Search (https://www.sanger.ac.uk/htgt/wge/find_crisprs)  Teslovich (http://csg.sph.umich.edu/willer/public/lipids2010/)  UCSC Genome Browser (https://genome.ucsc.edu)  VectorBase (https://www.vectorbase.org/)  VISTA (http://genome.lbl.gov/vista/index.shtml)  WormBase (https://www.wormbase.org)  Xenbase (http://www.xenbase.org/entry/)  ZFIN (https://zfin.org) |
| Ensembl Genomes | ArrayExpress  CATH  ChEMBL  ENA  Ensembl  IntAct  STRING  UniProt | BioModels (http://www.ebi.ac.uk/biomodels/)  BioSamples (https://www.ebi.ac.uk/biosamples/)  EFO (https://www.ebi.ac.uk/efo/)  EVA (https://www.ebi.ac.uk/eva/)  Expression Atlas (https://www.ebi.ac.uk/gxa/home)  FlyBase (https://flybase.org/)  GenBank (https://www.ncbi.nlm.nih.gov/genbank/)  GOA (https://www.ebi.ac.uk/GOA)  Gramene (http://www.gramene.org/)  HGNC (https://www.genenames.org/)  Identifiers.org (https://identifiers.org)  ImmunoDB(http://cegg.unige.ch/Insecta/immunod)  Joint Genome Institute (JGI)  KEGG (https://www.genome.jp/kegg/)  MEROPS (https://www.ebi.ac.uk/merops/)  mirBase (http://www.mirbase.org/)  NCBI (https://www.ncbi.nlm.nih.gov/)  OLS (https://www.ebi.ac.uk/ols/index)  Open Targets (https://www.opentargets.org/)  PHI-Base (http://www.phi-base.org/)  PomBase (https://www.pombase.org/)  Reactome (https://reactome.org)  Rfam (http://rfam.xfam.org/)  Rhea (https://www.rhea-db.org/)  RNAcentral (https://rnacentral.org)  SGD (https://www.yeastgenome.org/)  ToxoDB (https://toxodb.org/toxo/)  VectorBase (https://www.vectorbase.org/)  WormBase (https://www.wormbase.org) |
| Europe PMC | ArrayExpress  ChEBI  ChEMBL  EGA  ENA  Ensembl  Ensembl Genomes  IntAct  InterPro  PDBe  PRIDE  UniProt | Agricola (https://agricola.nal.usda.gov/)  BioModels (http://www.ebi.ac.uk/biomodels/)  BioProject (https://www.ncbi.nlm.nih.gov/bioproject)  BioStudies (https://www.ebi.ac.uk/biostudies/)  Cellosaurus (https://web.expasy.org/cellosaurus/)  Chinese Biological Abstracts (http://english.sibs.cas.cn/sp/CBA/CBADatabase/)  ClinicalTrials.gov (https://clinicaltrials.gov/)  Complex Portal (https://www.ebi.ac.uk/complexportal/home)  Crossref (https://www.crossref.org/)  dbSNP (https://www.ncbi.nlm.nih.gov/snp)  DisGeNET (http://www.disgenet.org/)  Dryad Digital Repository (https://datadryad.org/)  EBiSC (https://www.ebisc.org/)  EMDB (http://www.ebi.ac.uk/pdbe/emdb/)  EMPIAR (https://www.ebi.ac.uk/pdbe/emdb/empiar/)  EPO (https://www.epo.org/index.html)  EthOs Theses (British Library) (https://ethos.bl.uk/Home.do;j)  EudraCT (https://eudract.ema.europa.eu/)  FlyBase (https://flybase.org/)  Gene Ontology (http://geneontology.org/)  GenomeRNAi (http://genomernai.dkfz.de/)  GWAS Catalog (https://www.ebi.ac.uk/gwas/)  HGNC (https://www.genenames.org/)  HipSci (http:www.hipsci.org/)  Human DEPhOsphorylation Database (DEPOD) (https://www.depod.bioss.uni-freiburg.de/)  IGSR/1000 Genomes (https://www.internationalgenome.org)  Immune Epitope Database (http://www.iedb.org/home_v3.php)  iPTMnet (http://pir.georgetown.edu/iPTMnet)  MetaboLights (https://www.ebi.ac.uk/metabolights/)  MGnify (https://www.ebi.ac.uk/metagenomics/)  NeuroMorpho (http://neuromorpho.org/)  NHS Evidence (https://www.nice.org.uk/guidance)  OMIM (https://www.omim.org/)  Open Targets (https://www.opentargets.org/)  PANGAEA (https://www.pangaea.de/)  Pfam (https://pfam.xfam.org/)  PhenoMiner (http://boreas.mml.cam.ac.uk/phenominer/)  PubMed/MEDLINE NLM (https://www.ncbi.nlm.nih.gov/pubmed/)  Reactome (https://reactome.org)  RefSeq (https://www.ncbi.nlm.nih.gov/refseq/)  Rfam (https://rfam.xfam.org/)  RNAcentral (https://rnacentral.org)  Treefam (https://www.treefam.org/)  WikiPathways (https://www.wikipathways.org/)  WormBase (https://www.wormbase.org) |
| Human Protein Atlas | ArrayExpress  ChEMBL  Ensembl  Uniprot | Allen Brain Atlas (http://portal.brain-map.org)  Antibodypedia (https://www.antibodypedia.com)  Cellosaurus (https://web.expasy.org/cellosaurus/)  COSMIC (https://cancer.sanger.ac.uk/cosmic)  Drugbank (https://www.drugbank.ca/)  ENZYME (https://enzyme.expasy.org/)  Fantom5 (http://fantom.gsc.riken.jp/5/)  GTEx (https://gtexportal.org/home/)  Guide to Pharmacology (http://www.guidetopharmacology.org/)  KEGG (https://www.genome.jp/kegg/)  NCBI (https://www.ncbi.nlm.nih.gov/)  neXtprot (https://www.nextprot.org)  NucleaRDB (https://bio.tools/nucleardb)  PubMed (https://www.ncbi.nlm.nih.gov/pubmed/)  QuickGO (https://www.ebi.ac.uk/QuickGO/)  Reactome (https://reactome.org)  SNOMED CT (http://www.snomed.org)  TCDB (http://www.tcdb.org)  TFClass (https://omictools.com/tfclass-tool)  The Antibody Registry (http://antibodyregistry.org) |
| IntAct | CATH  ChEBI  ChEMBL  Ensembl  Ensembl Genomes  Europe PMC  InterPro  MINT  PRIDE  STRING  UniProt | APID (http://cicblade.dep.usal.es:8080/APID/init.action)  BioStudies (https://www.ebi.ac.uk/biostudies/)  Complex Portal (https://www.ebi.ac.uk/complexportal/home)  Cytoscape (https://www.cytoscape.org)  GeneMANIA (https://genemania.org)  HiPPIE (http://cbdm-01.zdv.uni-mainz.de/~mschaefer/hippie/information.php)  HPIDB (http://hpidb.igbb.msstate.edu)  I2D (https://www.accessdata.fda.gov/scripts/cder/iig/index.Cfm)  Identifiers.org (https://identifiers.org)  IID (https://www.accessdata.fda.gov/scripts/cder/iig/index.Cfm)  InnateDB (https://www.innatedb.com)  iRefIndex (http://irefindex.org/wiki/index.php?title=iRefIndex)  MatrixDB (http://matrixdb.univ-lyon1.fr)  Mentha (https://mentha.uniroma2.it/)  OLS (https://www.ebi.ac.uk/ols/index)  OmniPath (http://omnipathdb.org/)  Open Targets (https://www.opentargets.org/)  pathDIP (http://ophid.utoronto.ca/pathdip)  Reactome (https://reactome.org)  RNAcentral (https://rnacentral.org)  UniHI (https://omictools.com/unihi-tool) |
| InterPro | BRENDA  CATH  ChEMBL  ENA  Ensembl  Ensembl Genomes  Europe PMC  PDBe  UniProt | BioStudies (https://www.ebi.ac.uk/biostudies/)  CDD (https://www.ncbi.nlm.nih.gov/cdd)  Complex Portal (https://www.ebi.ac.uk/complexportal/home)  GOA (https://www.ebi.ac.uk/GOA)  HAMAP (https://hamap.expasy.org/)  HGNC (https://www.genenames.org/)  Identifiers.org (https://identifiers.org)  IntEnz (https://www.ebi.ac.uk/intenz/)  KEGG (https://www.genome.jp/kegg/)  MetaCyc (https://metacyc.org/)  MGnify (https://www.ebi.ac.uk/metagenomics/)  MobDB-Lite (http://protein.bio.unipd.it/mobidblite/)  OLS (https://www.ebi.ac.uk/ols/index)  Open Targets (https://www.opentargets.org/)  PANTHER (http://www.pantherdb.org)  Pfam (https://pfam.xfam.org/)  PIRSF (http://pir.georgetown.edu/pirwww/dbinfo/pirsf.shtml)Prosite  PRINTS (http://130.88.97.239/PRINTS/index.php)  ProDom (http://prodom.prabi.fr/prodom/current/html/home.php)  PROSITE (https://prosite.expasy.org)  Reactome (https://reactome.org)  SFLD (http://sfld.rbvi.ucsf.edu/django/)  SMART (http://smart.embl.de/)  SUPFAM (http://supfam.org)  TIGRFAMs (https://www.jcvi.org/tigrfams)  VectorBase (https://www.vectorbase.org)  WormBase (https://www.wormbase.org) |
| MINT | IntAct  PDBe  STRING  UniProt | PubMed (https://www.ncbi.nlm.nih.gov/pubmed/)  QuickGO (https://www.ebi.ac.uk/QuickGO/)  RCSB PDB (https://www.rcsb.org) |
| Orphadata | Ensembl  UniProt | HPO (https://hpo.jax.org/app/)  HGNC (https://www.genenames.org/)  LOVD (https://www.lovd.nl/)  Reactome (https://reactome.org/)  IUPHAR/BPS Guide to Pharmacology (https://www.guidetopharmacology.org/)  Genatlas (http://genatlas.medecine.univ-paris5.fr/)  ICD-10 (https://www.who.int/classifications/icd/icdonlineversions/en/)  ICD-11 (https://icd.who.int/en/)  OMIM (https://www.omim.org/)  GARD (https://rarediseases.info.nih.gov/)  UMLS (https://www.nlm.nih.gov/research/umls/index.html)  MeSH (https://www.ncbi.nlm.nih.gov/mesh)  MedDRA (https://www.meddra.org/) |
| PDBe | BRENDA  CATH  ChEBI  ChEMBL  ENA  Ensembl  Ensembl Genomes  Europe PMC  InterPro  MINT  STRING  UniProt | 3DComplex (http://shmoo.weizmann.ac.il/elevy/3dcomplexV6/Home.cgi)  BioStudies (https://www.ebi.ac.uk/biostudies/)  BMRB (http://www.bmrb.wisc.edu)  CCDC (https://www.ccdc.cam.ac.uk)  Complex Portal (https://www.ebi.ac.uk/complexportal/home)  Drugbank (https://www.drugbank.ca/)  EMDB (http://www.ebi.ac.uk/pdbe/emdb/)  EMPIAR (https://www.ebi.ac.uk/pdbe/emdb/empiar/)  Enzyme Portal (https://www.ebi.ac.uk/enzymeportal/)  GOA (https://www.ebi.ac.uk/GOA)  Identifiers.org (https://identifiers.org)  IntEnz (https://www.ebi.ac.uk/intenz/)  IRRMC (https://proteindiffraction.org)  Open Targets (https://www.opentargets.org/)  PDB_REDO (https://pdb-redo.eu)  PDBj (https://pdbj.org)  Pfam (https://pfam.xfam.org)  RCSB PDB (https://www.rcsb.org)  Reactome (https://reactome.org)  Rfam (http://rfam.xfam.org)  RNAcentral (https://rnacentral.org)  SASBDB (https://www.sasbdb.org)  SBGrid (https://sbgrid.org)  SCOP (http://scop.mrc-lmb.cam.ac.uk/scop/)  Taxonomy (https://www.ebi.ac.uk/ena/browse/taxonomy-service)  WormBase (https://www.wormbase.org) |
| PRIDE | ArrayExpress  EGA  ENA  Ensembl  Europe PMC  IntAct  UniProt | BioSamples (https://www.ebi.ac.uk/biosamples/)  BioStudies (https://www.ebi.ac.uk/biostudies/)  EFO (https://www.ebi.ac.uk/efo/)  Expression Atlas (https://www.ebi.ac.uk/gxa/home)  GPMDB (https://gpmdb.thegpm.org)  Identifiers.org (https://identifiers.org)  massIVE (https://massive.ucsd.edu/ProteoSAFe/static/massive.jsp)  MetaboLights (https://www.ebi.ac.uk/metabolights/)  MGnify (https://www.ebi.ac.uk/metagenomics/)  OLS (https://www.ebi.ac.uk/ols/index)  OmicsDI (https://www.omicsdi.org/)  Open Targets (https://www.opentargets.org/)  ProteomeCentral (http://proteomecentral.proteomexchange.org/cgi/GetDataset)  ProteomicsDB (https://www.proteomicsdb.org)  Reactome (https://reactome.org)  UCSC Genome Browser (https://genome.ucsc.edu)  WormBase (https://www.wormbase.org) |
| SILVA | ENA | Greengenes (http://greengenes.lbl.gov)  LPSN (http://www.bacterio.net/)  RDP (https://rdp.cme.msu.edu/)  RNAcentral (https://rnacentral.org) |
| STRING | Ensembl  Ensembl Genomes  IMEX Consortium  PDBe  Uniprot | BioCyc (https://biocyc.org/)  BioGrid (https://thebiogrid.org/)  COG (https://www.ncbi.nlm.nih.gov/COG/)  FlyBase (https://flybase.org/)  Gene Ontology (http://geneontology.org/)  KEGG (https://www.genome.jp/kegg/)  OMIM (https://www.omim.org/)  PubMed (https://www.ncbi.nlm.nih.gov/pubmed/)  Reactome (https://reactome.org)  RefSeq (https://www.ncbi.nlm.nih.gov/refseq/)  SIMAP (http://cube.univie.ac.at/resources/simap)  SwissModel (https://swissmodel.expasy.org/) |
| UniProt | ArrayExpress  BRENDA  CATH  ChEBI  ChEMBL  ENA  Ensembl  Ensembl Genomes  Europe PMC  Human Protein Atlas  IntAct  InterPro  MINT  Orphadata  PDBe  PRIDE  STRING | Allergome (http://www.allergome.org/)  ArachnoServer (http://www.arachnoserver.org)  Araport (https://www.araport.org/)  Bgee (https://bgee.org)  BindingDB (https://www.bindingdb.org/bind/index.jsp)  BioCyc (https://biocyc.org/)  BioGrid (https://thebiogrid.org/)  BioModels (http://www.ebi.ac.uk/biomodels/)  BioMuta (https://hive.biochemistry.gwu.edu/tools/biomuta/)  BioStudies (https://www.ebi.ac.uk/biostudies/)  CarbonylDB (http://digbio.missouri.edu/CarbonylDB/)  CAZy (http://www.cazy.org/)  CCDS (https://www.ncbi.nlm.nih.gov/CCDS)  CDD (https://www.ncbi.nlm.nih.gov/cdd)  CGD (http://www.candidagenome.org/)  ChiTaRS (http://chitars.md.biu.ac.il/)  CleanEx (https://cleanex.epfl.ch//)  ClinGen (https://www.clinicalgenome.org/)  ClinVar (https://www.ncbi.nlm.nih.gov/clinvar/)  CollecTF (http://www.collectf.org/)  Complex Portal (https://www.ebi.ac.uk/complexportal/home)  COMPLUYEAST-2DPAGE (http://compluyeast2dpage.dacya.ucm.es/)  ConoServer (http://www.conoserver.org/)  CORUM (http://mips.helmholtz-muenchen.de/corum/)  COSMIC (https://cancer.sanger.ac.uk/cosmic)  CTD (http://ctdbase.org/)  dbSNP (https://www.ncbi.nlm.nih.gov/snp)  DDBJ (https://www.ddbj.nig.ac.jp/)  Decipher (https://decipher.sanger.ac.uk/)  DEPOD (http://depod.bioss.uni-freiburg.de)  dictyBase (http://dictybase.org/)  DIP (https://dip.doe-mbi.ucla.edu/)  DisGeNET (http://www.disgenet.org/)  DisProt (http://disprot.bio.unipd.it)  DMDM (http://bioinf.umbc.edu/dmdm/)  DNASU (https://dnasu.org/DNASU/)  DOSAC-COBS-2DPAGE (http://www.dosac.unipa.it/2d/)  DrugBank (https://www.drugbank.ca/)  EchoBASE (https://www.york.ac.uk/res/thomas/)  EcoGene (http://www.ecogene.org/)  EFO (https://www.ebi.ac.uk/efo/)  eggNOG (http://eggnogdb.embl.de/)  ELM (http://elm.eu.org/)  EMPIAR (https://www.ebi.ac.uk/pdbe/emdb/empiar/)  ENZYME (https://enzyme.expasy.org/)  Enzyme Portal (https://www.ebi.ac.uk/enzymeportal/)  EPD (https://epd.epfl.ch//index.php)  ESP (http://evs.gs.washington.edu/EVS/)  ESTHER (http://bioweb.supagro.inra.fr/ESTHER/general?what=index)  euHCVdb (https://euhcvdb.ibcp.fr/euHCVdb/)  EuPathDB (https://eupathdb.org/eupathdb/)  EVA (https://www.ebi.ac.uk/eva/)  EvolutionaryTrace (http://lichtargelab.org/software/ETserver)  ExAC (http://exac.broadinstitute.org/)  Expression Atlas (https://www.ebi.ac.uk/gxa/home)  FlyBase (https://flybase.org/)  GenAtlas (http://genatlas.medecine.univ-paris5.fr/)  GenBank (https://www.ncbi.nlm.nih.gov/genbank/)  GeneCards (https://www.genecards.org/)  GeneDB (http://www.genedb.org/)  GeneID (https://www.ncbi.nlm.nih.gov/gene)  GeneReviews (https://www.ncbi.nlm.nih.gov/books/NBK1116)  Genevisible (https://genevisible.com/search)  GeneWiki (https://en.wikipedia.org/wiki/Portal:Gene_Wiki)  GenomeRNAi (http://genomernai.dkfz.de/)  GlyConnect (https://glyconnect.expasy.org)  GO (http://geneontology.org/)  GOA (https://www.ebi.ac.uk/GOA)  GPCRDB (http://gpcrdb.org)  Gramene (http://www.gramene.org/)  Guide to Pharmacology (http://www.guidetopharmacology.org/)  H-InvDB (http://www.h-invitational.jp/)  HAMAP (https://hamap.expasy.org/)  HGNC (https://www.genenames.org/)  HOGENOM (http://doua.prabi.fr/databases/hogenom/home.php)  HOVERGEN (http://pbil.univ-lyon1.fr/databases/hovergen.php)  HPA (https://www.proteinatlas.org/)  HUGE (http://www.kazusa.or.jp/huge/)  Identifiers.org (https://identifiers.org)  IGSR (http://www.internationalgenome.org/data-portal/sample)  IMGT_GENE-DB (http://www.imgt.org/genedb)  InParanoid (http://inparanoid.sbc.su.se/)  IntEnz (https://www.ebi.ac.uk/intenz/)  iPTMnet (http://pir.georgetown.edu/iPTMnet)  KEGG (https://www.genome.jp/kegg/)  KO (https://www.genome.jp/kegg/)  LegioList (http://genolist.pasteur.fr/LegioList/)  Leproma (ttps://mycobrowser.epfl.ch/)  MaizeGDB (https://www.maizegdb.org/)  MalaCards (https://www.malacards.org)  MaxQB (http://maxqb.biochem.mpg.de/mxdb/)  MEROPS (https://www.ebi.ac.uk/merops/)  MGI (http://www.informatics.jax.org/)  MGnify (https://www.ebi.ac.uk/metagenomics/)  Micado (http://genome.jouy.inra.fr/cgi-bin/micado/index.cgi)  MobiDB (http://mobidb.bio.unipd.it/)  ModBase (http://modbase.compbio.ucsf.edu/modbase-cgi/index.cgi)  MoonProt (http://moonlightingproteins.org/)  mycoCLAP (https://mycoclap.fungalgenomics.ca/mycoCLAP/)  neXtProt (https://www.nextprot.org)  OGP (http://usc_ogp_2ddatabase.cesga.es/cgi-bin/2d/2d.cgi)  OLS (https://www.ebi.ac.uk/ols/index)  OMA (https://omabrowser.org/)  OmicsDI (https://www.omicsdi.org/)  OMIM (http://www.omim.org/)  Open Targets (https://www.opentargets.org/)  Orphanet (https://www.orpha.net/)  OrthoDB (https://www.orthodb.org)  PANTHER (http://www.pantherdb.org)  PATRIC (https://patricbrc.org/)  PaxDb (https://pax-db.org)  PDBj (https://pdbj.org)  PeptideAtlas (http://www.peptideatlas.org)  PeroxiBase (http://peroxibase.toulouse.inra.fr/)  Pfam (https://pfam.xfam.org/)  PharmGKB (https://www.pharmgkb.org/)  PhosphoSitePlus (https://www.phosphosite.org)  PhylomeDB (http://phylomedb.org/)  PIR (http://pir.georgetown.edu/)  PIRSF (http://pir.georgetown.edu/pirwww/dbinfo/pirsf.shtml)  PMAP-CutDB (http://substrate.burnham.org/)  PomBase (https://www.pombase.org/)  PRINTS (http://130.88.97.239/PRINTS/index.php)  PRO (http://pir.georgetown.edu/pro/pro.shtml)  ProDom (http://prodom.prabi.fr/prodom/current/html/home.php)  ProMEX (http://promex.pph.univie.ac.at/promex/)  PROSITE (https://prosite.expasy.org)  ProtoNet (http://www.protonet.cs.huji.ac.il/)  PseudoCAP (http://www.pseudomonas.com/)  RCSB PDB (https://www.rcsb.org)  Reactome (https://reactome.org)  REBASE (http://rebase.neb.com/rebase/rebase.html)  RefSeq (https://www.ncbi.nlm.nih.gov/refseq/)  REPRODUCTION-2DPAGE (http://reprod.njmu.edu.cn/cgi-bin/2d/2d.cgi)  Rfam (http://rfam.xfam.org/)  RGD (https://rgd.mcw.edu/)  Rhea (https://www.rhea-db.org/)  Rouge (http://www.kazusa.or.jp/rouge/)  SABIO-RK (http://sabiork.h-its.org/)  SBKB (http://sbkb.org/)  SFLD (http://sfld.rbvi.ucsf.edu/django/)  SGD (https://www.yeastgenome.org/)  SignaLink (http://signalink.org/)  SIGNOR (https://signor.uniroma2.it/)  SMART (http://smart.embl.de/)  SMR (https://swissmodel.expasy.org/repository/)  SOURCE (http://source-search.princeton.edu/)  SUPFAM (http://supfam.org)  SWISS-2DPAGE (https://world-2dpage.expasy.org/swiss-2dpage/)  SWISS-MODEL (https://swissmodel.expasy.org/)  SwissLipids (http://www.swisslipids.org)  SwissPalm (https://swisspalm.org)  TAIR (https://www.arabidopsis.org/)  TCDB (http://www.tcdb.org)  TCGA (https://www.cancer.gov/about-nci/organization/ccg/research/structural-genomics/tcga)  TIGRFAMs (https://www.jcvi.org/tigrfams)  TubercuList (http://genolist.pasteur.fr/TubercuList/)  UCD-2DPAGE (https://proteomics-portal.ucd.ie/cgi-bin/2d/2d.cgi?spot=BRAIN_DLPFC_6-11:316&accession=P40926&data=all&database=human)  UCSC Genome Browser (https://genome.ucsc.edu)  UniCarbKB (http://www.unicarbkb.org)  UniGene (https://www.ncbi.nlm.nih.gov/unigene)  VectorBase (https://www.vectorbase.org/)  WormBase ParaSite (https://parasite.wormbase.org/index.html)  World-2DPAGE (https://world-2dpage.expasy.org/portal/)  WormBase (https://www.wormbase.org)  Xenbase (http://www.xenbase.org/entry/)  ZFIN (https://zfin.org) |

* The “**Links to additional data resources”** column enumerates 634 links outwards from this CDR set, for this March 2019 snapshot, to a total of 359 distinct data resources. Table S6 can be accessed in spreadsheet format in the Supporting Material Supporting.Material.CDR.infrastructure.12082019.xlsx file (DOI: 10.5281/zenodo.3468133 - https://zenodo.org/record/3468133), Tab “Table S6”.

Figure 4 includes data from the following Core Data Resources:

ArrayExpress, BRENDA, CATH, ChEBI, ChEMBL, EGA, ENA, Ensembl, Ensembl Genomes, EuropePMC, HPA, IntAct , InterPro, MINT, Orphadata, PDBe, PRIDE, SILVA, STRING, UniProt

**Figure 5. Heat map of the pairwise co-citation of the 12 ELIXIR Core Data Resources that are most frequently co-cited.**

The following steps were taken:

1-4. Steps 1-4 are the same as for Figure 3.

5. Having identified all CDR citations in terms of Resource name mentions, Resource accession number mentions and Key Article citations (recorded in the Fig3.Fig5.Step4.cdr_all tab in the Supporting Material [here](https://zenodo.org/record/3468133)), PMIDs that cited more than one CDR were used for a co-citation analysis. For each pairwise combination of CDRs, the number of common unique PMIDs were counted and displayed graphically in Figure 5 as the log of the co-citation count for each pair of resources.

**Table S7. Data from which Figure 5 is derived:**

| **Resource** | **Other resource** | **Log of**  **Co-citation**  **count** | **Resource** | **Other resource** | **Log of**  **Co-citation**  **count** |
| --- | --- | --- | --- | --- | --- |
| UniProt | The IMEx  Consortium | 2.47 | InterPro | UniProt | 2.86 |
| UniProt | STRING-db | 2.74 | InterPro | The IMEx  Consortium | 2.12 |
| UniProt | SILVA | 1.69 | InterPro | STRING-db | 2.35 |
| UniProt | PRIDE | 2.59 | InterPro | SILVA | 1.74 |
| UniProt | PDBe | 3.68 | InterPro | PRIDE | 1.96 |
| UniProt | InterPro | 2.86 | InterPro | PDBe | 2.66 |
| UniProt | Human Protein Atlas | 2.45 | InterPro | Human Protein Atlas | 1.63 |
| UniProt | Ensembl | 2.81 | InterPro | Ensembl | 2.31 |
| UniProt | ENA | 3.44 | InterPro | ENA | 2.82 |
| UniProt | CATH | 2.12 | InterPro | CATH | 1.96 |
| UniProt | ArrayExpress | 2.15 | InterPro | ArrayExpress | 1.95 |
| The IMEx Consortium | UniProt | 2.47 | Human Protein Atlas | UniProt | 2.45 |
| The IMEx Consortium | STRING-db | 2.70 | Human Protein Atlas | The IMEx  Consortium | 1.79 |
| The IMEx Consortium | SILVA | 0.30 | Human Protein Atlas | STRING-db | 2.21 |
| The IMEx Consortium | PRIDE | 1.79 | Human Protein Atlas | SILVA | 0.00 |
| The IMEx Consortium | PDBe | 2.09 | Human Protein Atlas | PRIDE | 2.09 |
| The IMEx Consortium | InterPro | 2.12 | Human Protein Atlas | PDBe | 2.309 |
| The IMEx Consortium | Human Protein Atlas | 1.79 | Human Protein Atlas | InterPro | 1.63 |
| The IMEx Consortium | Ensembl | 1.81 | Human Protein Atlas | Ensembl | 2.23 |
| The IMEx Consortium | ENA | 1.38 | Human Protein Atlas | ENA | 2.10 |
| The IMEx Consortium | CATH | 1.46 | Human Protein Atlas | CATH | 0.85 |
| The IMEx Consortium | ArrayExpress | 1.89 | Human Protein Atlas | ArrayExpress | 2.06 |
| STRING-db | UniProt | 2.74 | Ensembl | UniProt | 2.81 |
| STRING-db | The IMEx  Consortium | 2.70 | Ensembl | The IMEx  Consortium | 1.81 |
| STRING-db | SILVA | 1.00 | Ensembl | STRING-db | 2.18 |
| STRING-db | PRIDE | 2.38 | Ensembl | SILVA | 0.70 |
| STRING-db | PDBe | 2.41 | Ensembl | PRIDE | 1.97 |
| STRING-db | InterPro | 2.35 | Ensembl | PDBe | 2.46 |
| STRING-db | Human Protein Atlas | 2.21 | Ensembl | InterPro | 2.31 |
| STRING-db | Ensembl | 2.18 | Ensembl | Human Protein Atlas | 2.231 |
| STRING-db | ENA | 2.10 | Ensembl | ENA | 2.851 |
| STRING-db | CATH | 1.57 | Ensembl | CATH | 1.381 |
| STRING-db | ArrayExpress | 2.18 | Ensembl | ArrayExpress | 2.251 |
| SILVA | UniProt | 1.69 | ENA | UniProt | 3.44 |
| SILVA | The IMEx  Consortium | 0.30 | ENA | The IMEx  Consortium | 1.38 |
| SILVA | STRING-db | 1.00 | ENA | STRING-db | 2.10 |
| SILVA | PRIDE | 0.60 | ENA | SILVA | 2.78 |
| SILVA | PDBe | 1.26 | ENA | PRIDE | 2.03 |
| SILVA | InterPro | 1.74 | ENA | PDBe | 3.66 |
| SILVA | Human Protein Atlas | 0.00 | ENA | InterPro | 2.82 |
| SILVA | Ensembl | 0.70 | ENA | Human Protein Atlas | 2.10 |
| SILVA | ENA | 2.78 | ENA | Ensembl | 2.85 |
| SILVA | CATH | 0.60 | ENA | CATH | 1.46 |
| SILVA | ArrayExpress | 0.78 | ENA | ArrayExpress | 2.54 |
| PRIDE | UniProt | 2.59 | CATH | UniProt | 2.12 |
| PRIDE | The IMEx  Consortium | 1.79 | CATH | The IMEx  Consortium | 1.46 |
| PRIDE | STRING-db | 2.38 | CATH | STRING-db | 1.57 |
| PRIDE | SILVA | 0.60 | CATH | SILVA | 0.60 |
| PRIDE | PDBe | 2.05 | CATH | PRIDE | 0.70 |
| PRIDE | InterPro | 1.96 | CATH | PDBe | 2.52 |
| PRIDE | Human Protein Atlas | 2.09 | CATH | InterPro | 1.96 |
| PRIDE | Ensembl | 1.97 | CATH | Human Protein Atlas | 0.85 |
| PRIDE | ENA | 2.03 | CATH | Ensembl | 1.38 |
| PRIDE | CATH | 0.70 | CATH | ENA | 1.46 |
| PRIDE | ArrayExpress | 1.96 | CATH | ArrayExpress | 0.85 |
| PDBe | UniProt | 3.68 | ArrayExpress | UniProt | 2.15 |
| PDBe | The IMEx  Consortium | 2.09 | ArrayExpress | The IMEx  Consortium | 1.89 |
| PDBe | STRING-db | 2.41 | ArrayExpress | STRING-db | 2.18 |
| PDBe | SILVA | 1.26 | ArrayExpress | SILVA | 0.78 |
| PDBe | PRIDE | 2.05 | ArrayExpress | PRIDE | 1.96 |
| PDBe | InterPro | 2.66 | ArrayExpress | PDBe | 1.85 |
| PDBe | Human Protein Atlas | 2.30 | ArrayExpress | InterPro | 1.95 |
| PDBe | Ensembl | 2.46 | ArrayExpress | Human Protein Atlas | 2.06 |
| PDBe | ENA | 3.66 | ArrayExpress | Ensembl | 2.25 |
| PDBe | CATH | 2.52 | ArrayExpress | ENA | 2.54 |
| PDBe | ArrayExpress | 1.85 | ArrayExpress | CATH | 0.85 |

Figure 5 includes data from the following Core Data Resources:

ArrayExpress, CATH, ENA, Ensembl, HPA, IntAct/MINT, InterPro, PDBe, PRIDE, SILVA, STRING, UniProt. Co-citations do occur across the full set of CDRs, but the less frequently occurring of these were removed for legibility of Figure 5.

**Figure 6. Horizon of assured funding**

In December 2018, the Core Data Resource managers were sent a survey, asking:

Q1: How many Full Time Employees (FTEs) do you have committed funding for [Resource Name] on 1 January in the years below?”

The years for which data was requested were 2019-2024, inclusive.

**Table S8. Data from which Figure 6 is derived:**

Data for years 2014-2018 are as for Figure 1.

| **Year** | **2019** | **2020** | **2021** | **2022** | **2023** | **2024** |
| --- | --- | --- | --- | --- | --- | --- |
| **Assured Full Time Positions** | 346.9 | 306.0 | 250.0 | 115.2 | 63.0 | 39.0 |

Figure 6 includes data from the following Core Data Resources:

ArrayExpress, BRENDA, CATH, ChEBI, ChEMBL, EGA, ENA, Ensembl, Ensembl Genomes

EuropePMC, HPA, IntAct/MINT, InterPro, Orphadata, PDBe, PRIDE, SILVA, STRING, UniProt
